# Just add water: Urban blue spaces increase avian richness and functional diversity

**DOI:** 10.1101/2025.06.18.660350

**Authors:** Matthew C. Morgan, Rodney Forster, Charlotte R. Hopkins, Africa Gómez

## Abstract

Urban blue spaces are highly valuable for both people and nature, providing key ecosystem services, including flood alleviation, pollution absorption, microclimate regulation, benefits to human health and wellbeing, and habitat provision. Crucially, urban blue spaces support biodiversity, including threatened species, and despite often being small, may have disproportionate effects on their surrounding environment, acting as critical habitats within urban systems. However, research on the role of urban blue spaces within ecological contexts remains limited. Here, we assessed urban bird communities across green and blue spaces to quantify the ecological effects of urban water bodies. We surveyed birds along 22 paired 1 km transects in the city of Kingston Upon Hull, UK, recording species and abundance across both winter and breeding seasons. Our findings indicate that blue spaces significantly increase bird species richness during summer (*P* = 0.016), though not in winter. However, we found that the taxonomic distinctiveness of bird communities is consistently greater around blue spaces across both seasons (*P* < 0.05). Similarly, functional diversity based on species-level ecological traits was more varied around water (*P* = 0.01). In addition, we show that urban blue spaces could be important for avian conservation, supporting more red and amber-listed species than green spaces during the summer (*P* = < 0.05). Overall, our results show that urban blue spaces play a critical ecological role within cities by enhancing the complexity of avian communities, which in turn could improve human wellbeing and contribute to urban sustainability.

## Introduction

The process of urbanisation is increasing globally, caused by growing human populations and rural-urban migration (Yehya et al., 2024). Urban areas now host over 56% of the world’s population, predicted to increase to 68% by 2050 (United Nations: Department of Economic and Social Affairs: Population Division, 2019). Urban expansion alters land use and land cover, often leading to habitat loss and biodiversity declines (Li et al., 2022). Remnants of natural land cover in urban settings, such as rivers and woodlands, can act as a refuge for nature, but are usually degraded and fragmented (Di Giulio et al., 2009; Sanetra et al., 2024). Similarly, artificial semi-natural environments, such as parks, lakes and gardens, can have positive effects on urban biodiversity (Hill et al., 2017; Delahay et al., 2023), providing suitable habitats for species declining in rural areas, such as Hedgehogs (*Erinaceus europaeus*) (Williams et al., 2015). Collectively, these natural and semi-natural urban environments, referred to as urban blue and green spaces, are recognised for their benefits to human health and well-being (Lauers, 2021), as well as their ecological value (Knight et al., 2022). Blue and green spaces provide a range of ecosystem and social benefits, including flood prevention (Krivtsov et al., 2020), environmental cooling (Yu et al., 2020), air purification (Vieira et al., 2018), and provide opportunities for recreation (Börger et al., 2021) and direct interactions with nature (Fleming & Shwartz, 2023; Stanford et al., 2024). Within this context, research has primarily focused on green spaces, such as amenity grasslands, parks and street trees, resulting in a broad acceptance of their environmental and social value (Reyes-Riveros et al., 2021; Jabbar et al., 2022). In contrast, blue spaces, defined as all natural and manmade surface waters in urban environments (Grellier et al., 2017; Smith et al., 2021), including rivers, canals and park lakes, are less well studied (White et al., 2020) despite some evidence supporting comparable social and ecological value (Brückner et al., 2021).

Wetland habitats have declined globally by 23% since 1700 (Fluet-Chouinard et al., 2023), a proportion of which can be directly attributed to the process of urbanisation (Burgin et al., 2016; Ballut-Dajud et al., 2022). For example, infilling, culverting and land drainage, which reduce blue space cover, are all common practices during urban development (McEwen et al., 2020; Thornhill et al., 2022). In addition, any remaining water courses are often heavily modified through the process of channelisation, removing meandering bends and soft sloping edges, and polluted with urban runoff (Wang et al., 2001) and wastewater (Silva et al., 2024). Despite all this, urban blue spaces can provide therapeutic and restorative environments for people (Finlay et al., 2015; Subiza-Pérez et al., 2020; Smith et al., 2022), and access to them can improve mental health and well-being (McDougall et al., 2022), promote physical exercise (Tan et al., 2021), and increase socialisation (Smith et al., 2022). They also provide important habitat for biodiversity, providing links across urban landscapes via blue-green corridors (e.g., rivers and drains) (Hyseni et al., 2021) and stepping stones for migratory wetland species (Krivtsov et al., 2022).

Wetlands support a disproportionately large amount of biodiversity by area (Dudgeon et al., 2006) and can influence the environment beyond their footprint through aquatic subsidies (Lewis-Phillips et al., 2020) and emergent pollinating insects (Murail et al., 2024). Around 25% of freshwater fauna is threatened with extinction (Sayer et al., 2025), and freshwater urban blue spaces have the potential to play a role in reversing this trend. For example, urban water bodies in towns and cities have been shown to act as refuges for populations of endangered Water Voles (*Arvicola amphibius*) (Brzeziński et al., 2018; Leivesley et al., 2021). Common wetland species found across urban areas are also important, providing ecosystem services which directly improve living conditions for humans. Riparian plants, such as Common Reeds (*Phragmites australis*), remove chemical pollutants from urban run-off (Swanson et al., 2017; Wu et al., 2023), insectivorous birds, such as Hirundines, help regulate insect populations (Roseo et al., 2024), and charismatic species, such as the Mute Swan (*Cygnus olor*), provide opportunities for human-nature interactions (Duke & Soulsbury, 2021). Although urban blue spaces have demonstrated benefits for biodiversity, their ecological value in urban settings remains understudied, particularly amid rising urbanisation and the concurrent biodiversity and climate crises. Research around urban ecology has focused mainly on biodiversity across the urban-rural gradients (McKinney, 2008; Husby et al., 2020; Wang et al., 2020), and the size, connectivity and quality of blue-green spaces (Norton et al., 2016; Hyseni et al., 2021), with few studies disentangling distinctions between green and blue spaces.

In this study, we quantified the influence of urban blue spaces on biodiversity using birds as ecological indicators. Fieldwork was conducted in Kingston Upon Hull, a densely populated major UK city, where we surveyed urban areas with and without blue spaces, using paired transects. We recorded bird species and abundance, which was combined with ecological trait data and conservation status, to assess both taxonomic and functional diversity, as well as conservation value.

## Methods

### Study Site

Kingston upon Hull (53.7675° N, 0.3273° W) is a major UK city situated on the northern bank of the Humber Estuary (Figure 1). The area is highly urbanised with 80% of land cover being impervious surfaces, 15% green space and 5% blue space. The city has a population of 267,000 people and is ranked as the fourth most deprived local authority in England (Hull City Council, 2025). There is a dense urban core surrounded on the east, north, and west by agricultural land, with extensive networks of open and culverted drains, which converge into three major waterways as they enter the city. The River Hull, running centrally, is a tidal system connected to the Humber Estuary, which delineates the southern limit of the city. To the east of the River Hull lies the Beverley and Barmston Drain, and to the west, the Holderness Drain. Several inland water bodies are also located within city parks, including a 1 km–long boating lake. Due to low-lying land, the region is susceptible to flooding (Coulthard & Frostick, 2010), and several forms of blue-green infrastructure have been integrated into the city to manage water, including ‘aqua greens’ and sustainable drainage systems (Hull City Council, 2015; Environment Agency, 2024).

**Figure 1.**
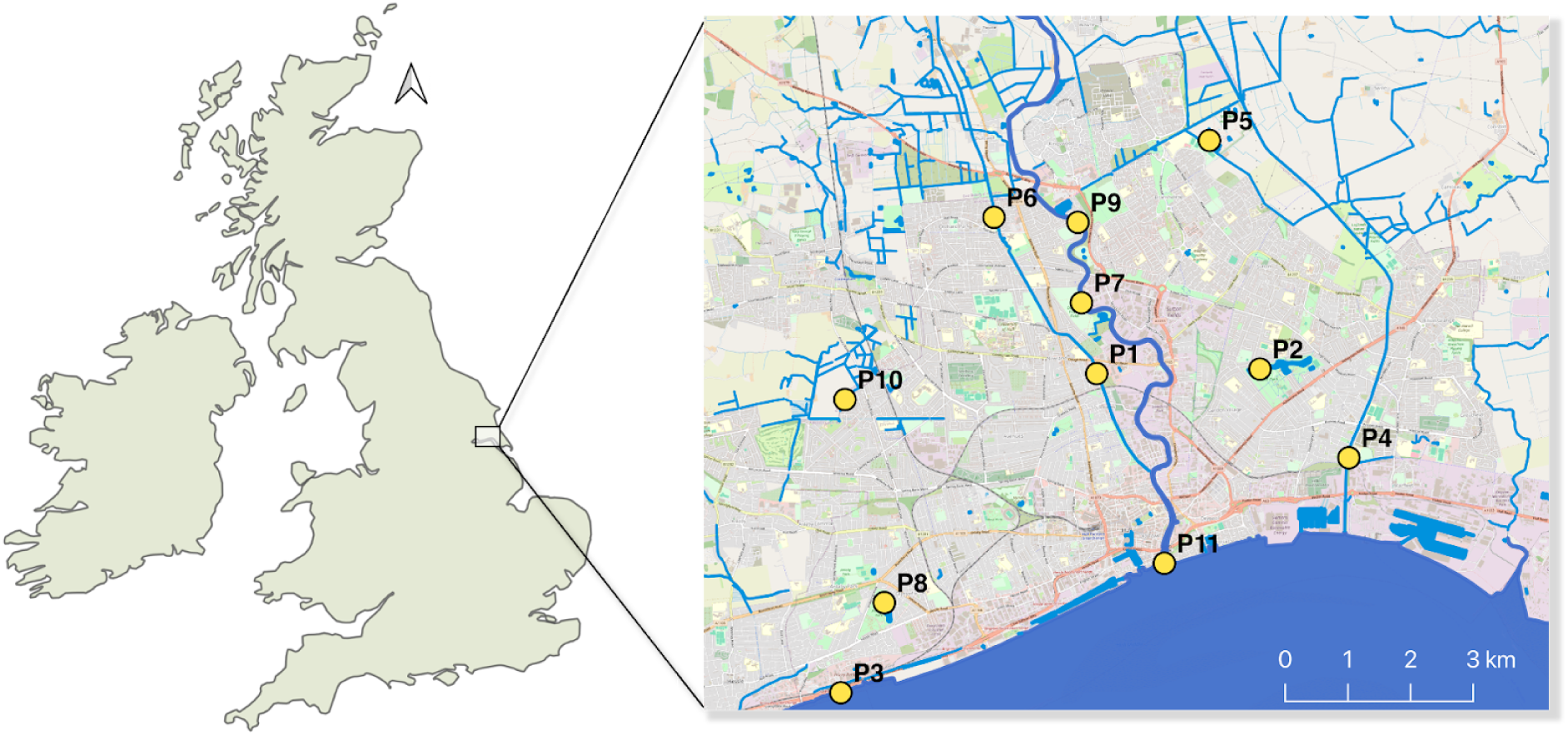
Map of study site location. Map showing the United Kingdom (left) and the location of Kingston Upon Hull (right). The distribution of the main blue spaces in the city are marked in blue. The location of transect pairs are shown with yellow dots.

### Data Collection

We used paired transects (n = 22), one-kilometre long by 200 m wide, (one with water, and one without water), in the main, following the British Trust for Ornithology (BTO) methodology for Breeding Bird Surveys (BBS) and Winter Bird Surveys (WBS) (BTO, 2018). Transect routes were designed using a combination of satellite imagery, local knowledge and public accessibility (see Figure 2 for details). Transect pairs were located as close as possible to one another without overlap, while maintaining comparable land cover, other than the presence of a significant blue space (e.g., river, lake, drain, reservoir, estuary). Distances between transects varied due to the fragmented and irregular distribution of natural spaces across the city, and deviations from straight lines occurred due to inaccessible land or physical obstructions (e.g., fences, overgrown paths). In transects with water, the observer walked alongside the water body along a public footpath if available, or as close as accessibility allowed. Each transect was divided into five 200 m subsections to facilitate data collection, and three horizontal distance bands: 0–25 m, 25–100 m, and 100+ m. All birds encountered on the transects were recorded to species level using visual and acoustic cues, along with distance band and abundance. Birds in flight were also noted. Observers used the Merlin Bird ID Application developed by Cornell University as an additional aid in the confirmation and detection of birds during the transects. In total, 88 km were surveyed across four recording periods, by two experienced ornithologists: twice in the late summer (June 2023-2024), once in winter (January 2024), and once in early summer (May 2024). During late summer surveys, only adult birds were recorded. Paired transects were surveyed on the same day by the same observer, within a 2-hour window, starting at 7:00 am in the summer and 09:00 am in the winter. Each pair of transects was carried out by the same observer for all four recording periods. Observers used binoculars to identify birds and mobile phones to navigate transects using Google My Maps, where the transects were plotted. To minimise observer bias and standardise methods, a pilot survey of two full transects was conducted by both observers before official surveys began.

**Figure 2.**
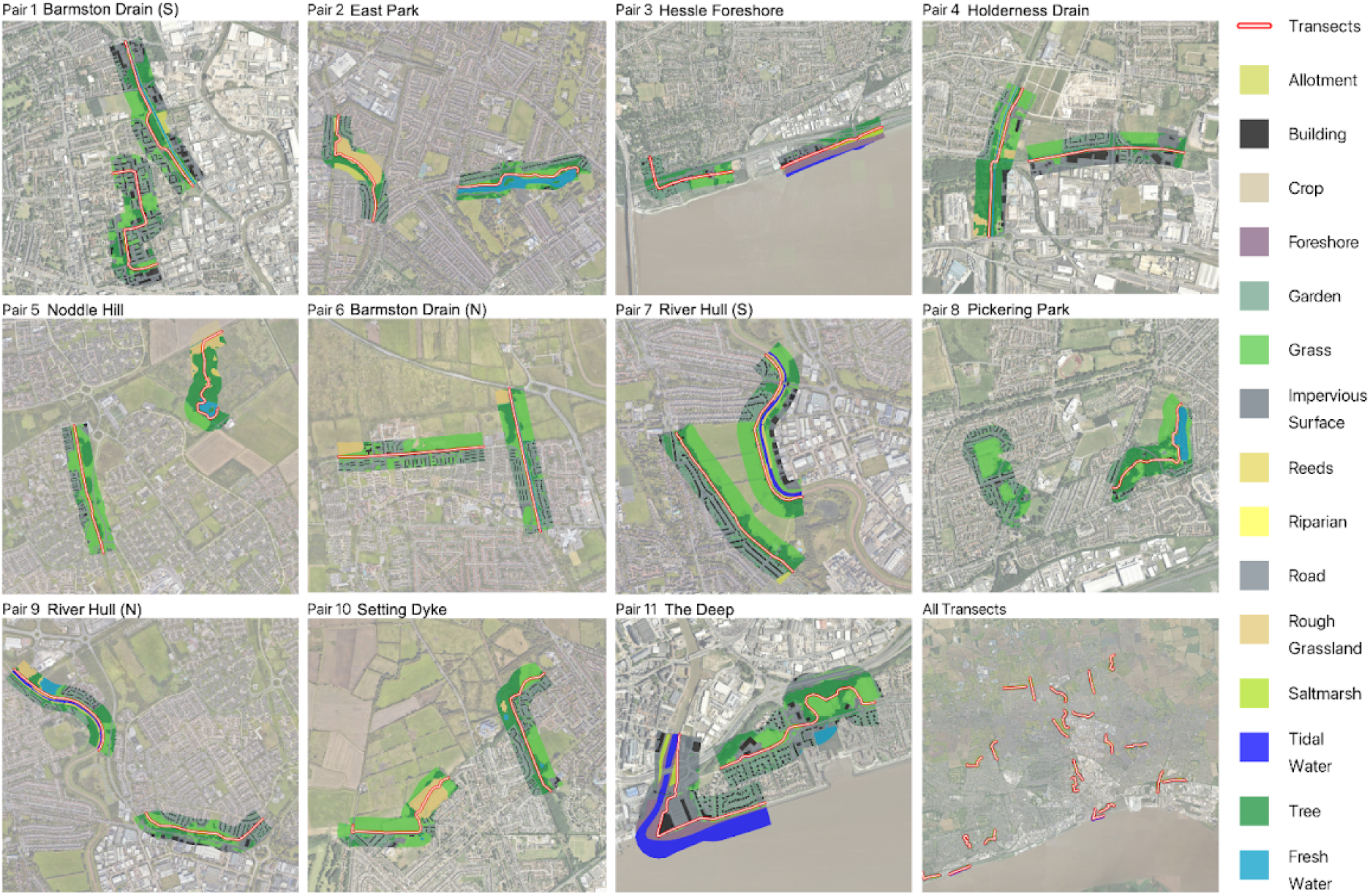
Detailed maps of all transect pairs. Classified land cover at ≤ 10 m resolution across all transects (n = 22). See Supplementary Table 2 for transect descriptions.

Land cover across transects was quantified using a combination of land use data (Ordnance Survey, 2023), European Space Agency WorldCover map 2021 (10 m resolution) (Zanaga et al., 2022) and manual digitisation (see Figure 2). Ordnance Survey (OS) land use vector layers were used to account for *Buildings*, *Gardens*, *Roads* (line data buffered by 5 m), *Amenity Grasslands*, *Surface Water*, *Tidal Water*, *Foreshore*, and *Crops*. The remaining gaps were then filled with the WorldCover data, providing cover for *Trees*, *Impervious Surfaces*, and *Grasslands* (where missing), without replacing OS layers. Manual digitisation, informed by Google Satellite imagery and ground truthing, was used to add *Rough Grassland*, *Riparian Vegetation* and *Saltmarsh*. Land cover types with low occurrence and uneven spatial distribution were grouped for analysis (see Supplementary Table 1).

### Data Analysis

All data analyses were conducted in R version 4.2.2 (R Core Team, 2025) within RStudio version 2025.05.1 (RStudio Team, 2025), using base R functions and additional packages. Data manipulation and transformation were performed using the *tidyverse* package (Wickham et al., 2019). Species accumulation curves were generated with the *iNEXT* package (Chao et al., 2014; Hsieh et al., 2024). Alpha and beta diversity metrics were calculated using the *vegan* package (Oksanen et al., 2024). Detection probability estimates were modelled with the *Distance* package (Miller & Clarke-Wolf, 2023). Functional diversity metrics were assessed using the *SYNCSA* package (Debastiani & Pillar, 2012). Figures were created using *ggplot2* (Wickham, 2016) and formatted with *gridExtra* (Auguie, 2017).

Before analysing bird records, sampling effort across transects was assessed with species accumulation curves to check sufficient asymptotes had been achieved for transects with and without water (see Supplementary Fig. 1). Next, total species richness and abundance data from the three summer sampling periods were pooled and averaged into a single summer dataset. This was done after confirming there were no significant differences across periods using a repeated measures ANOVA, treating transect as a random effect (species richness: Df = 2, F value = 0.121, p-value = 0.886; bird abundance: Df = 2, F value = 6.66, p-value = 0.849; see Supplementary Table 3). Finally, land cover across transect pairs was compared using Wilcoxon-signed rank tests (WSRT), which confirmed the largest difference in land cover between transect pairs was blue space (W = 66, Z = −3.30, adj. p-value = 0.007), see Supplementary Fig. 2 for further details.

For both summer and winter datasets, raw and adjusted bird count data per transect were analysed using biodiversity metrics (Shannon’s Index, Simpson’s Index, Pielou’s evenness, and Species Richness), which were compared between transects with and without water. Raw count data included pooled counts from all distance bands and birds in flight per transect. Adjusted count data were corrected using detection probability estimates (DPE). DPE were modelled using the *Distance* package (Miller and Clarke-Wolf, 2023) to account for any effects of distance, bird size, transect visibility, water presence, and observer on detectability. Only data from the fixed distance bands were included (0-25 m and 25-100 m) as recommended by Buckland *et al*. (2001). Data for species-specific models was limited, so bird species were aggregated into three categories based on bird mass (g) extracted from AvoNet (Tobias et al., 2022), small (< 35 g), medium (36-400 g) and large (> 400g). Visibility of each transect subsection (n = 440) was categorised based on the observer’s line of sight. Visibility categories are as follows: open transects had largely unobstructed lines of sight on both sides, of around 50–100 m with minimal obstructions, semi-open transects had variable visibility, generally providing at least 25 m; in some cases, one side of the transect was closed while the other remained open, closed transects had restricted lines of sight on both horizontal and vertical planes, of less than 25 m along most of the section. Detection decay over distance was modelled using Half-normal detection functions as the distance data was binomial (see Supplementary Fig. 3 and Supplementary Table 4). Results were then compared across paired conditions (transects with and without blue space), using WSRT.

Unadjusted raw count data were used for the rest of the analysis to retain aerial species (e.g., Hirundines) and all distance bands. Community dissimilarities (beta diversity) between transects with and without water were assessed using Bray-Curtis dissimilarity (SIMPER). Fisher’s Exact tests were then carried out on non-aquatic species identified as significant with SIMPER.

Taxonomic distinctiveness and functional diversity of bird communities in each transect were assessed using four taxonomic levels (species, genus, family, order) and 33 unique species-level trait combinations based on body size, trophic niche, and primary lifestyle extracted from AvoNet (Tobias et al., 2022) (see Supplementary Table 5 and 6 for details). Average taxonomic distinctiveness (avTD) was calculated from a taxonomic dissimilarity matrix generated using the *tax2dist* function from the *vegan* package (Oksanen et al., 2022). Functional diversity (RaoQ) and functional redundancy were computed using the rao.diversity function from the *SYNCSA* package (Debastiani & Pillar, 2012). Differences in taxonomic and functional diversity between transects with and without water were tested using WSRT. Community-level differences in functional traits were tested using the PERMANOVA, applied to a Gower dissimilarity matrix based on species-level trait data. Differences in trait dispersion between transects with and without water were assessed using the betadisper function.

Total bird biomass was calculated by multiplying species mass by count. Non-aquatic biomass was calculated by excluding species assigned as *Wetland* by Avonet (see supplementary Table 5). Bird biomass per transect was log transformed and compared using WSRT across both seasons.

The number of species with red, amber, and green conservation status as listed in the Birds of Conservation Concern 5 (BoCC5) (Stanbury et al., 2021) was compared across transects with and without blue space for both seasons, using presence-absence data, tested with WSRT. Introduced species, Canada goose *(Branta canadensis*), Ring-necked parakeet (*Psittacula krameri*), and Pheasant (*Phasianus colchicus*), which lacked BoCC5 status, were treated as green, owing to their IUCN status of Least Concern (see supplementary Table 5).

## Results

In total, 14,708 birds were counted, totalling 79 species, 36 families and 15 orders, across 88 km of transects and four survey periods. Species accumulation curves confirmed that sampling effort was sufficient to reveal abundance and species richness patterns in transects with and without water, across both seasons. In summer and winter, on average, transects with water had 24% more species, and bird abundance was 60% and 56% higher, respectively (see supplementary Fig. 1). The most frequently recorded species were Woodpigeon (*Columba palumbus*) n = 713, Carrion crow (*Corvus corone*) n = 372, Blackbird (*Turdus merula*) n = 360 and Wren (*Troglodytes troglodytes*) n = 300 (see Fig. 3a). The most frequently recorded wetland specialists were Mallard (*Anas platyrhynchos*) ranked 19th and Moorhen (*Gallinula chloropus*) ranked 24th (see Fig. 3a). The five most abundant species were Woodpigeon (n = 2053), Greylag geese (*Anser anser*) n = 1435, Rock dove (*Columba livia*) n = 966, Starling (*Sturnus vulgaris*) n = 913, and Carrion crow n = 877 (see Fig. 3b). The top 25 most abundant species made up 90% of all records. Notable species with single records included Marsh harrier (*Circus aeruginosus*), Water rail (*Rallus aquaticus*) and Grey wagtail (*Motacilla cinerea*), see Supplementary Table 5 for further details.

**Figure 3.**
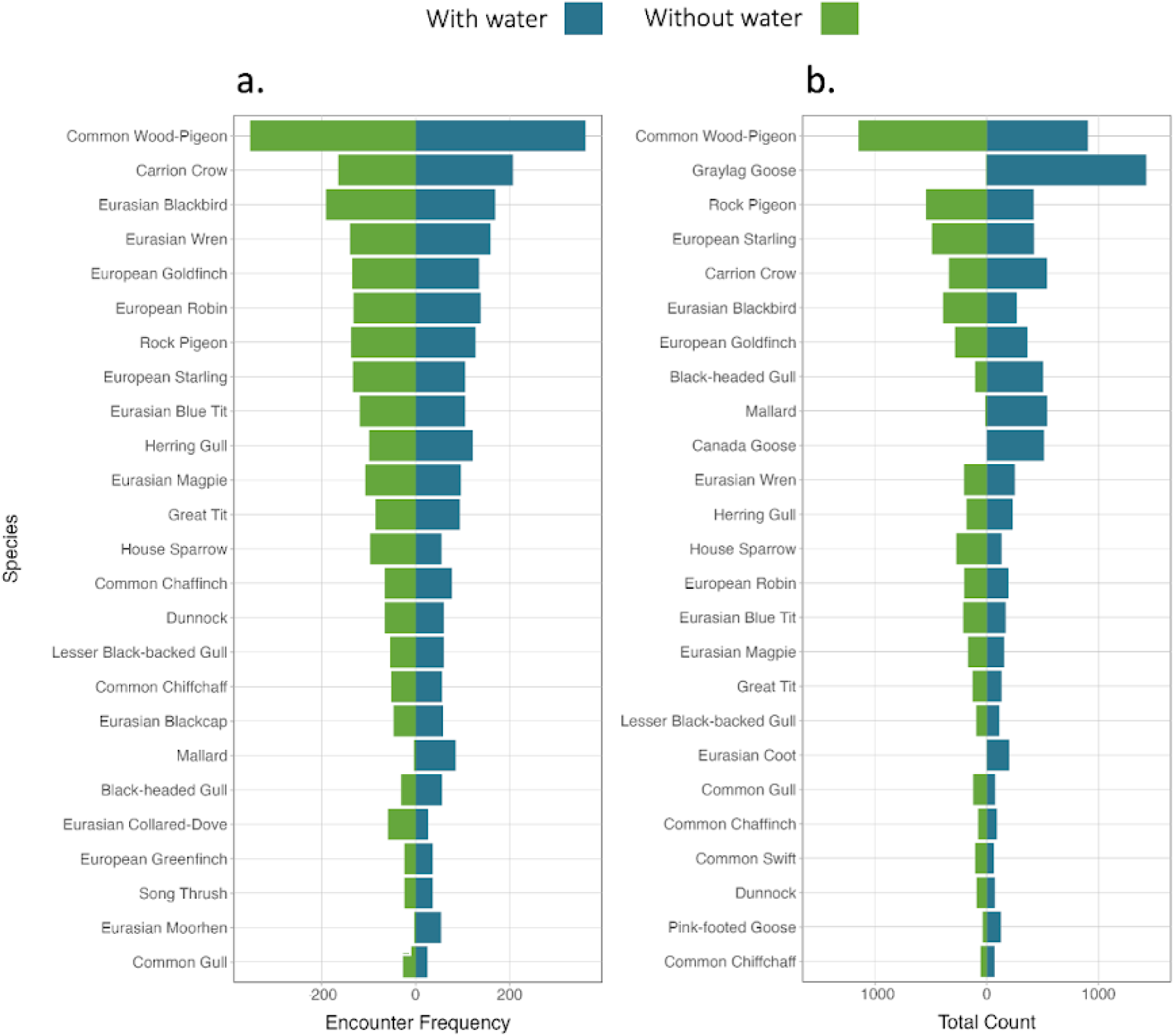
Encounter frequency and abundance of birds. Ranked diagrams showing the top 25 most frequently recorded bird species (a.) and the most abundant bird species (b.) across all transects with and without water (x-axis).

Out of the 79 bird species recorded, 27 species were only present in the transects with water, consisting of 20 wetland specialists and 7 non-aquatic species. Transects without water had no unique species. After testing with WSRT, total bird species richness was significantly higher in transects with water in the summer season for both raw counts (V = 51.5, p-value = 0.016) and DPE-adjusted counts (V = 64, p-value = 0.007). Species richness was not significantly higher in transects with water in the winter. No statistical differences between transects with and without water were found using Shannon’s, Simpson’s or Pielou’s evenness. Average taxonomic distinctiveness was significantly higher in transects with water in both the summer (V = 57, p-value = 0.032) and winter (V = 65, p-value = 0.002) (see Fig. 4), but no differences were found with RaoQ functional diversity or redundancy. See Supplementary Table 7 for a summary of all taxonomic and functional diversity analyses.

**Figure 4.**
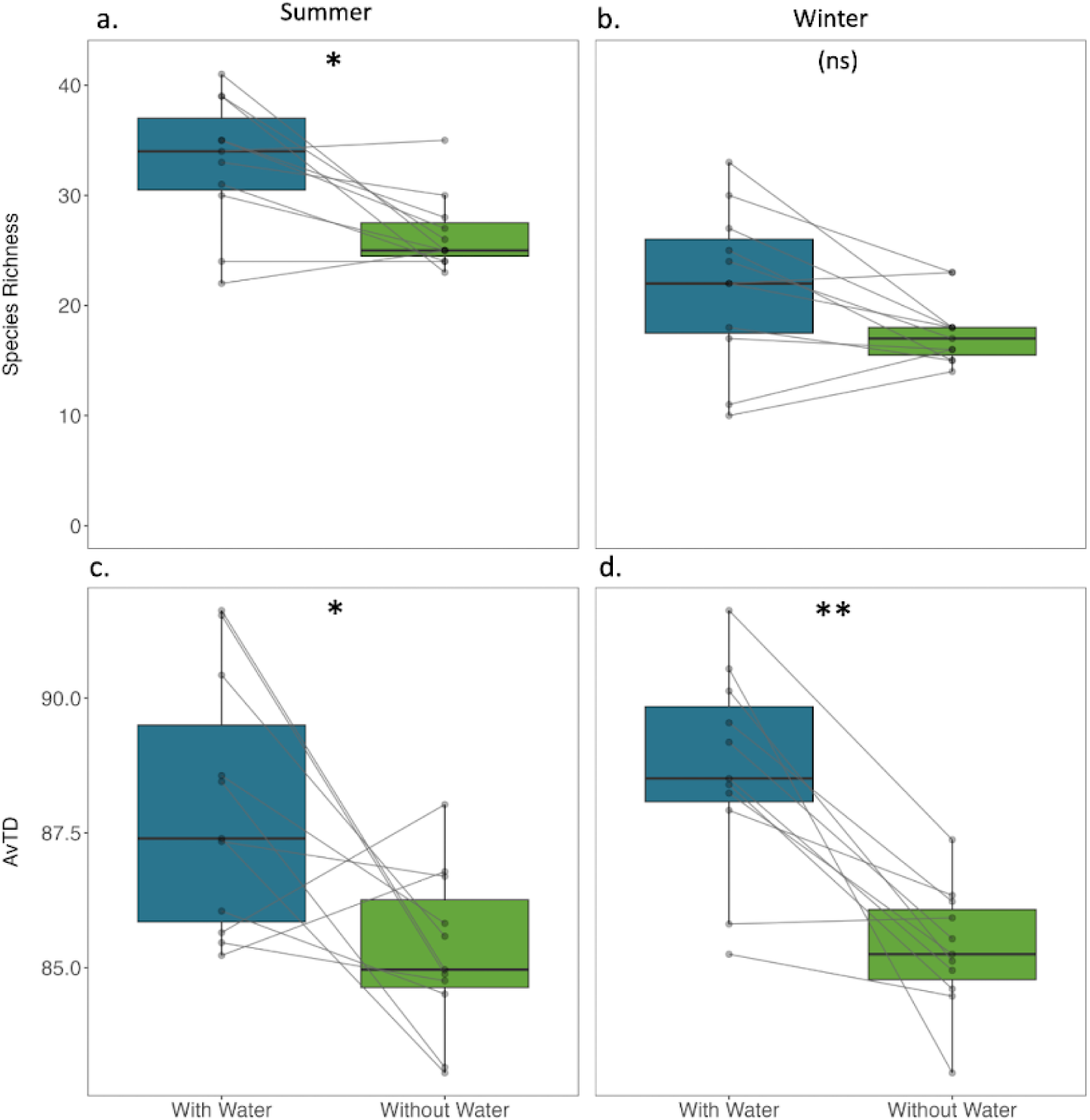
Comparison of biodiversity metrics. Total species richness and Average Taxonomic Distinctiveness (AvTD) in transects with and without water during the summer (left) and winter (right) surveys. Lines represent pairwise comparisons (Wilcoxon signed-rank test). Significant differences are indicated by * (p < 0.05) and ** (p < 0.001).

In the summer, SIMPER analysis identified 13 species that significantly contributed to community dissimilarities between transects, due to their higher abundances in transects with water. Five of these species were non-aquatic (i.e., not wetland specialists), Linnet (*Carduelis cannabina*), Greater Whitethroat (*Sylvia communis*), Willow warbler (*Phylloscopus trochilus*) and Stock dove (*Columba oenas*), but only Swallow (*Hirundo rustica*) remained statistically significant after testing presence-absence across transects with Fisher’s exact test (p-value = 0.009). No species was more abundant in the transects without water in the summer. In the winter, nine species were identified with SIMPER as being significantly different across transect conditions, three of which were non-aquatic. Redwing (*Turdus Iliacus*) was more abundant in transects with water, whereas Blackbird (*Turdus merula*) and Collared-Dove (*Streptopelia decaocto*) were more abundant in transects without water. However, none of these species remained statistically significant following Fisher’s exact test; see Table 1 for further details.

**Table 1:**
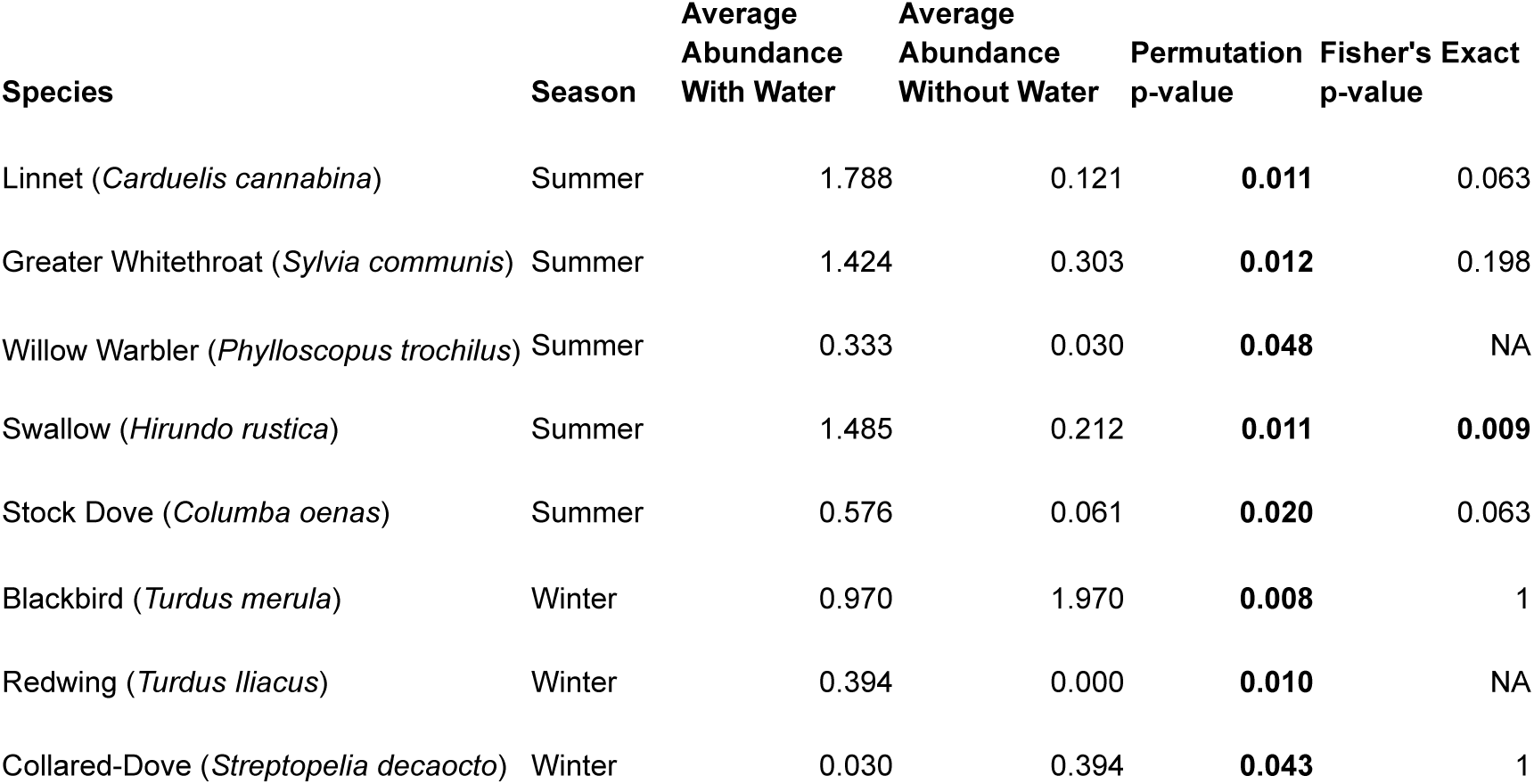
Summary table for SIMPER analysis and Fisher’s Exact tests. Results of non-aquatic species which had significant abundance dissimilarities between transects with and without water are shown. Columns 3-5 represent SIMPER outputs, the permutation p-value represents the probability that a species’ contribution to dissimilarity under random groupings equals or exceeds the observed contribution. Column 6 represents Fisher’s Exact test p-value, note that this test was only performed when the species had been recorded at a minimum of 5 transects.

| Species | Season | Average<br>Abundance<br>With Water | Average<br>Abundance<br>Without Water | Permutation<br>p-value | Fisher's Exact<br>p-value |
| --- | --- | --- | --- | --- | --- |
| Linnet (*Carduelis cannabina*) | Summer | 1.788 | 0.121 | **0.011** | 0.063 |
| Greater Whitethroat (*Sylvia communis*) | Summer | 1.424 | 0.303 | **0.012** | 0.198 |
| Willow Warbler (*Phylloscopus trochilus*) | Summer | 0.333 | 0.030 | **0.048** | NA |
| Swallow (*Hirundo rustica*) | Summer | 1.485 | 0.212 | **0.011** | **0.009** |
| Stock Dove (*Columba oenas*) | Summer | 0.576 | 0.061 | **0.020** | 0.063 |
| Blackbird (*Turdus merula*) | Winter | 0.970 | 1.970 | **0.008** | 1 |
| Redwing (*Turdus Iliacus*) | Winter | 0.394 | 0.000 | **0.010** | NA |
| Collared-Dove (*Streptopelia decaocto*) | Winter | 0.030 | 0.394 | **0.043** | 1 |

A Principal Coordinates Analysis (PCoA) based on Gower dissimilarity of species-level trait data, revealed distinct communities between transects with and without water, tested with a PERMANOVA (R² = 0.122, F = 2.79, p = 0.01) (See Fig. 5). Bird communities across transects with water were associatied with high body mass, herbivory, and predatory traits, while transects without water were associated with generalists, with granivorous and omnivorous traits. Multivariate dispersion analysis also showed that there is a significant difference in within-group variability (F = 15.76, p < 0.001), showing that transects with water had significantly higher functional dispersion (mean difference = –0.097, p < 0.001).

**Figure 5.**
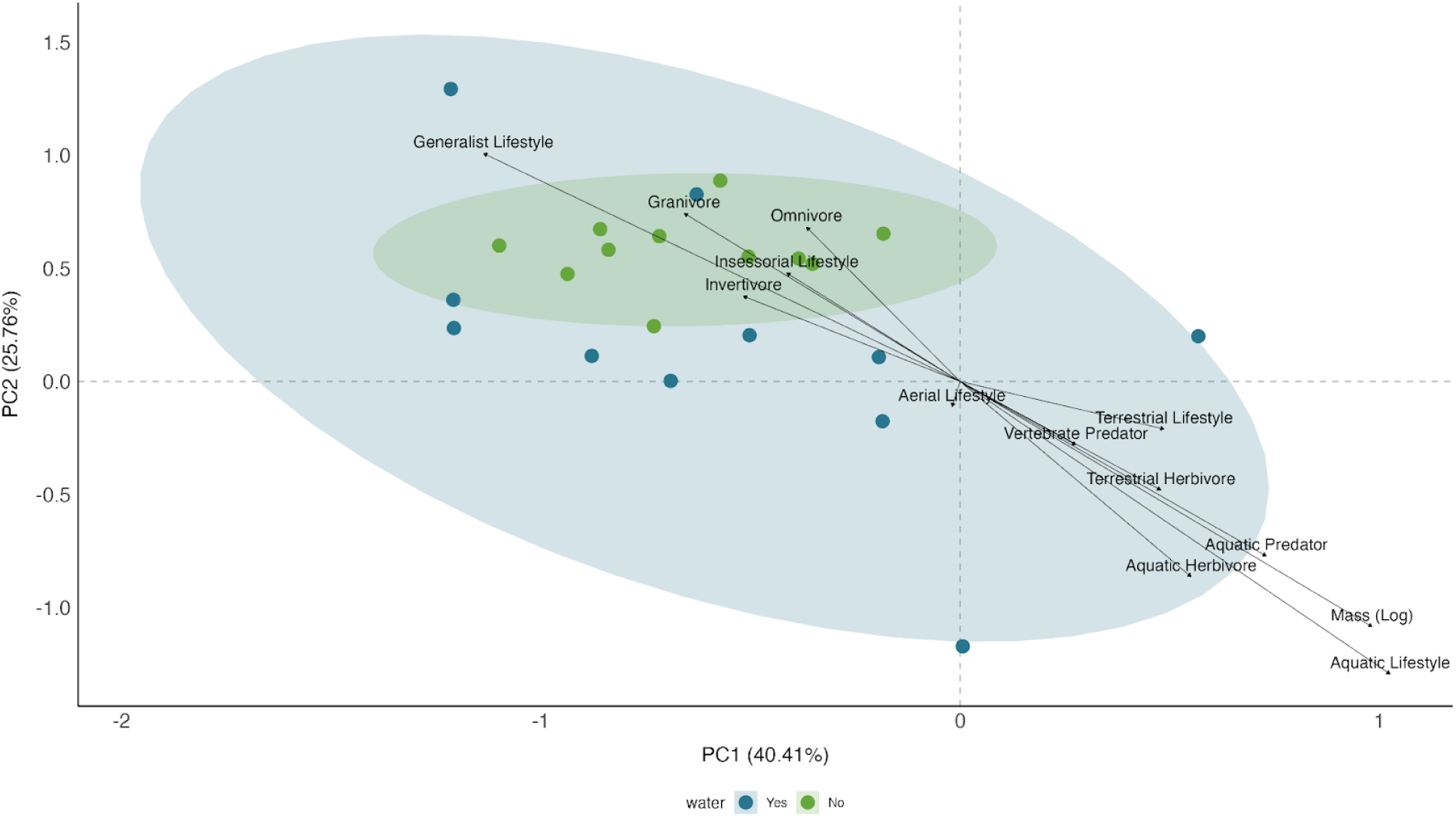
Principal Coordinates Analysis (PCoA) of functional traits. PCoA illustrating functional trait composition of bird communities from pooled seasons in transects with and without water. Points represent species-level trait combinations per transect, with ellipses indicating 95% confidence intervals around centroids for transects with and without water. Axes represent variation in pairwise community dissimilarity (Bray-Curtis). Arrows indicate the magnitude and direction of functional traits in the dataset.

In both summer and winter, 9 out of the 11 paired transects had higher bird biomass in the transects with water. In summer, transects with water had an average biomass of 218 kg, more than five times that of transects without water (44 kg). Variability was also higher in transects with water (Min: 17 kg, Max: 1177 kg, SD: 383 kg) than in those without (Min: 8 kg, Max: 40 kg, SD: 9 kg). In winter, average biomass in transects with water was 191 kg, compared to 58 kg in those without (a threefold difference), and variability remained greater in transects with water (Min: 23 kg, Max: 511 kg, SD: 187 kg) than in those without (Min: 28 kg, Max: 161 kg, SD: 36 kg). WSRT revealed statistically significant differences in bird biomass between transects with and without water in both summer (*V* = 61, *p* = 0.010) and winter (*V* = 59, *p* = 0.019). However, when aquatic species were removed, the differences of were no longer significant (see Supplementary Table 8 and Supplementary Fig. 4).

More than 50% of the species recorded in the study were species of conservation concern according to the BoCC5 assessment, with 13 species (16.5%) Red-listed, and 27 species (34.2%) Amber-listed. When species were aggregated by conservation status, transects with water had significantly more Amber-listed (V = 50.5, p-value = 0.022) and Red-listed (V = 42, p-value = 0.021) species in the summer, but only significantly more Amber-listed species in the winter (V = 60, p-value = 0.017), see Fig. 6. In the summer, transects with water supported four unique Amber-listed species, Reed bunting (*Emberiza schoeniclus*), Gadwall (*Mareca strepera*), Grey Wagtail (*Motacilla cinerea*) and Reed Warbler (*Acrocephalus schoenobaenus*), and a single Red-listed species, Mistle Thrush (*Turdus viscivorus*). In the winter, transects with water supported five unique Amber-listed species, notably Marsh Harrier (*Circus aeruginosus*), Redshank (*Tringa totanus*) and Wigeon (*Mareca penelope*), and three Red-listed species, Lapwing (*Vanellus vanellus*), Curlew (*Numenius arquata*) and Pochard (*Aythya ferina*).

**Figure 6.**
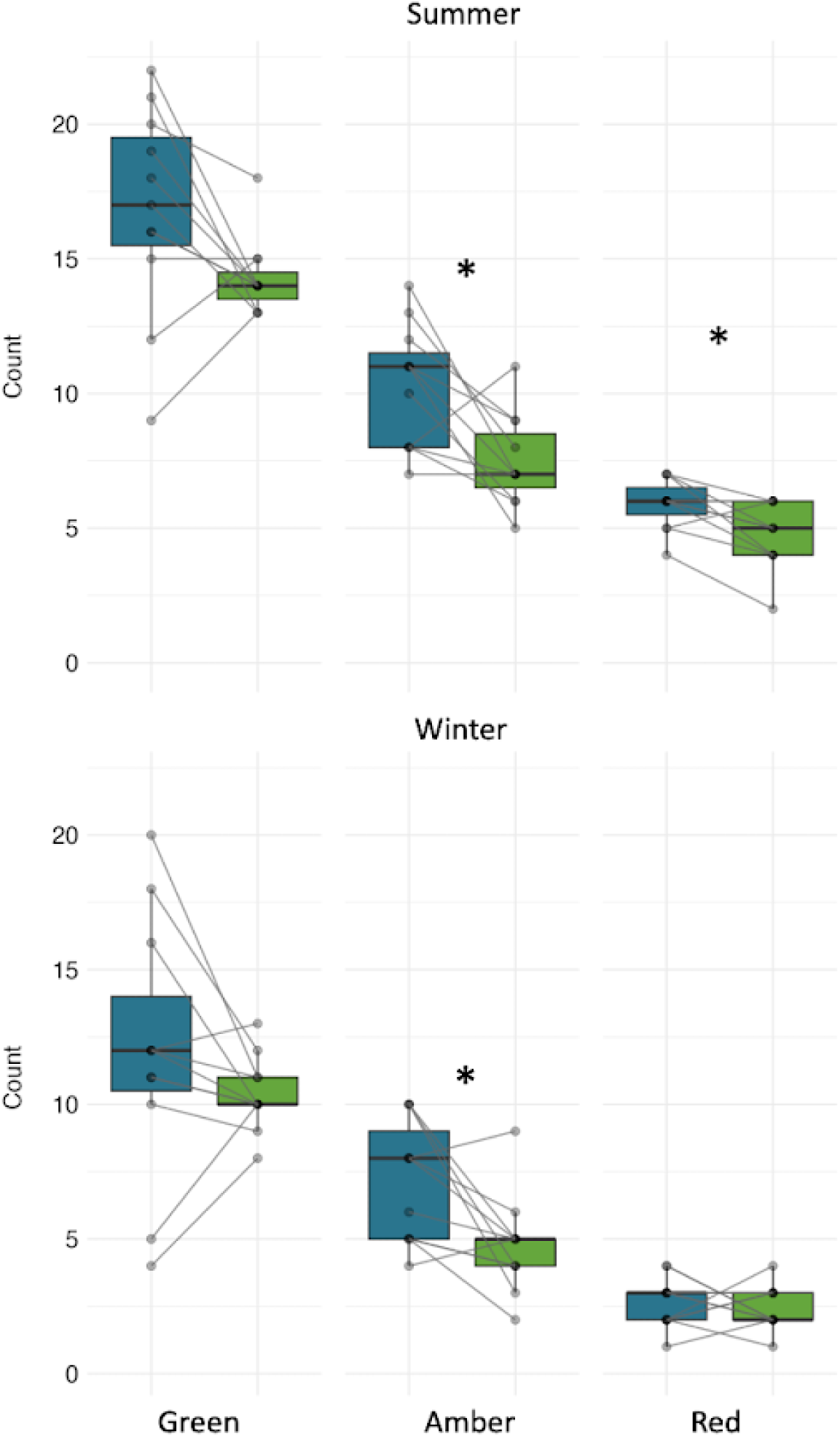
Comparison of conservation priority species. Number of bird species aggregated by conservation status (BoCC5) in summer and winter transects with and without water. Lines represent pairwise comparisons (Wilcoxon signed-rank test). Significant results are indicated by * (p < 0.05). See Supplementary Table 9 for full results.

## Discussion

By examining both taxonomic and functional aspects of urban bird communities, we have shown that blue spaces support greater bird species richness, abundance, and biomass compared to similar urban habitats without water bodies. Moreover, bird communities associated with blue spaces were more taxonomically distinct and exhibited a wider range of ecological traits, suggesting that urban wetlands may contribute to greater ecosystem resilience. In addition, we also found that blue spaces support more biomass and conservation priority species, underlining their important role within urban ecology and conservation.

### Taxonomic diversity

Our findings show that urban areas with water (blue space) support greater bird diversity than those without, with 27 bird species occurring only in transects with blue space. Urban parks with lakes were the most diverse habitat surveyed, supporting both aquatic and non-aquatic bird communities through a combination of habitats, including open grassland, mature trees, and ornamental planting, which border marginal aquatic habitats such as *Iris sp*. and *Baraganium sp.,* creating an ecotone between terrestrial and aquatic systems. In addition, their larger and deeper water bodies could also support diving birds such as Tufted Duck (*Aythya fuliga*), Great Crested Grebe (*Podiceps cristatus*) and Goosander (*Mergus merganser*) which were absent from shallow blue spaces, such as dykes and drains.

The capacity for urban wetlands to support greater bird diversity than comparable habitats has been demonstrated in other studies, including a coastal-inland comparison from Chile, which found coastal urban areas had higher species richness by supporting both inland and coastal bird communities (Graells et al., 2022). Our results also align with other urban studies, which demonstrate a general positive effect of water on avian species richness (Barbosa et al., 2020; Huang et al., 2022; Xie et al., 2022). Some studies also show urban wetlands can support more bird abundance and species richness than rural wetlands (McKinney et al., 2011), although this is not always the case (Aubrechtová et al., 2023). Here, we show that the positive relationship between urban blue spaces and bird species richness was strongest in the summer breeding season, potentially linked to summer migrants, such as Reed warbler (*Acrocephalus scirpaceus*) and House martins (*Delichon urbicum*), which benefit from emergent aquatic insects which are absent in the winter. Although other migratory species, such as the Redwing (*Turdus iliacus*), overwinter in the UK, our findings suggest they have less association with water bodies. This result is consistent with previous research, which found that farmland ponds have less activity from winter migrants than summer migrants (Lewis-Phillips et al., 2019).

The lack of blue space effect in winter could also be associated with environmental differences between the summer and winter months. Wetland birds are adapted to dynamic systems and have flexible behavioural responses, including foraging, to adapt to environmental change (Beerens et al., 2011). As the UK is prone to seasonal winter flooding (Soriano-Redondo et al., 2016) and our case study was conducted in a low-lying region, wetland bird species may have been less concentrated around the permanent blue spaces due to birds exploiting different seasonal resources. For example, we found wetland birds such as Moorhen (*Gallinula chloropus*) and Coot (*Fulica atra*) utilising flooded playing fields, and mixed species flocks of geese were observed grazing away from permanent water bodies.

Transects with blue spaces had greater abundance and species richness overall, but alpha biodiversity indices were not significantly different across transects with and without water. The similarity across these metrics could be explained by common synanthropic bird species found in abundance across the study area. In total, 90% of the total bird abundance was composed of 25 species, and the top 25 most frequently recorded birds made up 88% of all records. This contrasts with findings from other urban bird studies, where just 3–5 species made up 65–90% of recorded individuals (Peris & Montelongo, 2014; Thompson et al., 2022). However, despite the increased diversity of common birds in our study, it is likely that these common species still masked the effects of wetland specialists, which were encountered less frequently and often in smaller numbers.

### Functional diversity

Bird communities around blue spaces had significantly higher taxonomic distinctiveness, an indicator of phylogenetic diversity, than those around green spaces. As phylogenetic diversity of birds has been shown to decline with urban intensification (Santos et al., 2024), as in plants (Knapp et al., 2017), this suggests blue spaces may play an important role in preserving phylogenetically distinct communities within cities. We also found that bird communities around blue spaces had significantly higher variation of ecological traits, likely associated with the broad ecological range of blue spaces in the study, which included estuaries, tidal rivers, freshwater drains, lakes, and reservoirs. For example, aquatic predators such as Kingfisher (*Alcedo atthis*) and Little Grebe (*Tachybaptus ruficollis*) were unique to transects with water, as were some terrestrial herbivores, such as Canada Geese (*Branta canadensis*). These findings, which go beyond the perspectives of taxon counting (Miller et al., 2018), demonstrate that the additional niches provided by urban wetland habitats, not only increase the number of species using the area, but also the heterogeneity of functional traits, essential for ecosystem resilience (Gladstone-Gallagher et al., 2019). In addition, we found that bird biomass was significantly higher in transects with water, due to the effect of wetland species, which often have large bodies, associated with thermoregulation and buoyancy (Gutiérrez, 2014). Large wetland birds, such as Grey Heron (*Ardea cinerea*), also contribute to broader ecological functionality. By using both terrestrial and aquatic resources, they can perform important ecological roles, such as acting as vectors for plants and aquatic invertebrates (Navarro-Ramos et al., 2022).

### Aquatic subsidies

Along with the 27 species only being recorded within transects with water, five non-aquatic species were encountered more frequently and in greater abundance across transects with blue spaces. This suggests that non-aquatic bird species are benefiting from aquatic subsidies (e.g., emergent insects), which are smaller in magnitude than terrestrial-aquatic subsidies, such as tree leaves falling in a pond (Earl & Semlitsch, 2013), but higher in nutritional quality and energy density (Schindler & Smits, 2017). Here, we found that insectivorous Swallows (*Hirundo rustica*) have a strong association with urban blue spaces, and evidence that granivorous species such as Linnet (*Carduelis cannabina*) can also benefit. The benefits of aquatic subsidies have already been documented in farmland ponds, where Lewis-Phillips et al. (2020) found that ponds act as “insect chimneys”, allowing increased densities of local bird communities. However, the effects of urban water bodies on avian productivity remain underexplored compared to other urban resources. For example, garden bird-feeders have been shown to increase community evenness amongst garden birds (Plummer et al., 2019), and anthropogenic food items are known to increase the productivity of wetland species, such as Wood Stork (*Mycteria americana*) (Evans & Gawlik, 2020). Aquatic-terrestrial nutrient transfer from urban ponds, drains and rivers is seldom acknowledged, but as we have shown here, water bodies in cities are likely benefiting both wetland specialists and non-wetland species. When considering their relatively small area compared to green spaces, and their capacity to support increased species richness and abundance, along with evidence of resource subsidy benefits, urban blue spaces may function as critical habitats for birds. Much like keystone species, these systems could be considered “keystone habitats”, which have disproportionately large impacts on biodiversity relative to their size (Hitchman et al., 2018).

### Conservation significance and human benefits of bird diversity

We have shown that urban bird communities can be highly diverse and not necessarily made up of common species. Of the 79 bird species recorded in this study, 13 are listed as Red and 27 as Amber in The Birds of Conservation Concern 5 (BoCC5), a UK-based population status assessment (Stanbury et al., 2021). In general, blue spaces had more priority species, as well as priority species unique to those areas, such as Reed-bunting (*Emberiza schoeniclus*) and Lapwing (*Vanellus vanellus*). These findings align with previous studies showing the value of urban areas for avian conservation (Exantus et al., 2025), in particular those which have emphasised the importance of urban wetlands (Yang et al., 2025). Therefore, in cities, where space is at a premium, provisioning, protecting and establishing wetlands is important for bird communities. Furthermore, compensating for wetland loss with other habitat types is likely to result in reduced conservation value.

Although birds of conservation concern are present within cities, especially around water bodies (Yang et al., 2025), urban areas are not an ideal replacement for natural systems. Artificial conditions in urban areas can have negative impacts on bird fitness, for instance, the popular activities of feeding garden birds and ‘feeding the ducks” (Chapman & Jones, 2009) can increase disease transmission and cause deformities due to nutritional imbalances (Coogan et al., 2018), such as angel wings in swans (Arican et al., 2019). Other examples include reduced reproductive success, associated with low food availability and quality (Seress et al., 2020), and phenotypic changes, such as reduced eye size associated with light pollution (Hahs et al., 2023; Jones et al., 2023). Therefore, although urban ecosystems can support declining species, these unexpected refuges can have damaging long-term effects. However, most urban adaptation studies have focused on passerines, while adaptations of wetland species remain understudied (Minias, 2016), despite them showing a strong phylogenetic signal towards urban tolerance (Callaghan et al., 2019). Some wetland birds can have increased reproductive performance in urban areas, due to reduced predation, shown with Coots (*Fulica atra*) in Poland (Minias, 2016), and anthropogenic food items, shown with Wood Storks (*Mycteria americana*) in South Florida (Evans & Gawlik, 2020). Therefore, despite the drawbacks of urban areas, in the absence of better alternatives, they are likely providing important space for nature, particularly wetland birds.

### Human-Nature Interactions

Encounters with wildlife in blue-green places can strengthen human-nature connectedness (Bell et al., 2018; Folmer et al., 2019) and both perceived and actual species richness can enhance well-being (Dallimer et al., 2012). The increased abundance and biomass of birds around urban blue spaces, especially the presence of large-bodied wetland specialists, suggests blue spaces could facilitate exposure and accessibility to wildlife more effectively than urban habitats without water. Charismatic wetland species, such as the Mute Swan (*Cygnus olor*), among the UK’s most favourite birds (Lindo, 2015), are especially likely to capture public interest. Supporting this, Duke and Soulsbury (2021) found that waterfowl made up the majority of human-wildlife interactions in a study of urban blue spaces. In addition, exposure to bird song has been linked to improved well-being (Ratcliffe, 2021) and the perceived restorative capacity of environments (Fisher et al., 2021). Therefore, the higher bird abundance and diversity of blue spaces could afford greater therapeutic qualities than other urban spaces. These effects would be strongest in the summer, when summer migrants including Reed Warbler (*Acrocephalus scirpaceus*) and Sedge Warbler (*Acrocephalus schoenobaenus*) call abundantly. As green social prescribing has proven successful in the UK and momentum in the field is expected to continue (Wadsworth et al., 2024), health practitioners should consider how blue and green spaces differ from an ecological perspective, and how the therapeutic and restorative benefits of blue-green spaces could be influenced by fine scale differences in biodiversity. As shown here, bird biodiversity and biomass in cities can be concentrated across blue spaces, making them viable options for nature-based health interventions and activities.

### Rewilding for people and nature

To achieve urban sustainability, urban planners are challenged with improving the quality, size and connectivity of natural spaces (Beninde et al., 2015), and reducing light (Aguilera & González, 2023), noise (Jeon et al., 2010), and air pollution (Zhang et al., 2022). Yet, urban environments are highly complex socio-ecological systems, where space is at a premium and the value of accessible nature is often outweighed by other forms of urban growth, such as commercial development and urban sprawl (Hu et al., 2024). Furthermore, the provision and improvement of natural spaces can also be costly and are often unrealistic in less affluent areas. However, as wetlands support disproportionate amounts of biodiversity by area, urban blue spaces, if managed correctly, can offer highly effective options for rewilding cities. For example, Garrowby Orchard, included in this study, showed a huge positive ecological response after a culverted water course was reopened (‘daylighted’). The new habitat was quickly recolonised by fish and amphibians, attracting Little Egret (Egretta garzetta), Grey Heron (Ardea cinerea) and Kingfisher (Alcedo atthis). Another example included here is Noddle Hill, a site which rewilded itself after abandonment (passive rewilding), which has developed distinct bird communities when compared to adjacent farmland (Broughton et al., 2022), and its expansive reedbed ecosystems now support Marsh Harriers (*Circus aeruginosus*). Located on the edge of the city, these sites benefit from low light pollution and strong ecological connectivity with the surrounding landscape, whilst remaining easily accessible to people. Elsewhere, canal edges are being softened with coir rolls to improve habitat quality (Stoddart, 2017), man-made rafts in urban marinas are providing nesting ground for Common Terns (Sterna hirundo) (Simms, 2021), and vertical arrays of artificial rock pools on seawalls provide habitat for intertidal fauna (Bone et al., 2024). By reinstating blue spaces and improving the management of existing permanent wetland habitats (e.g., urban rivers, canals, dykes) through a reconciliation approach, balancing conservation and water management (Tölgyesi et al., 2022), urban blue spaces can support biodiversity gains which align with sustainable urban development targets.

## Conclusion

Using birds as indicators, we have shown that urban wetlands possess significant ecological and social value by examining both taxonomic and functional aspects of urban bird communities. Our findings show that bird communities associated with urban blue spaces exhibit greater species richness, abundance, and biomass compared to areas lacking blue spaces. Furthermore, these communities are more taxonomically distinct than those in green spaces and encompass a wide range of ecological traits, which may enhance the resilience of urban ecosystems. Importantly, we also discovered that urban blue spaces support more conservation priority species than comparable habitats. These ecological benefits are likely to translate into human benefits through increased exposure to nature, highlighting the role of blue spaces in promoting both biodiversity and human well-being. Together, these findings provide critical insights for advancing urban sustainability and guiding conservation efforts in increasingly urbanised landscapes.

## Supporting information

See supplementary material here.

## Acknowledgements

We gratefully acknowledge the University of Hull for funding this research. We also thank Hull City Council, the Yorkshire Wildlife Trust, and the Environment Agency for their encouragement and support of this work. Special thanks go to the University of Hull Rewild Cluster, particularly Clare Cowgill, for their ongoing assistance.

## Supplementary data

Supplemetary data can be found here **10.5281/zenodo.15641790**.

## Data availability

All raw data and R code used to produce results in this study are available on [github](https://github.com/MCMorgan06/Birds_Blue_Spaces) and zonodo **10.5281/zenodo.15641790** to promote transparency and reproducibility.

