## Supplementary material for "Just add water: Urban blue spaces increase avian richness and functional diversity": See supplementary material here.

### Supplementary Materials

Supplementary Table 1: Overview of land cover groupings. Original land cover classifications and respective groupings used for analysis.

| Original Land Covers | Land Cover Grouping for Analysis |
|---|---|
| Fresh Water, Reeds, Salt Marsh, Foreshore, Riparian, Tidal Water | Blue Space |
| Garden, Allotment, Crops | Garden |
| Grass, Rough Grass | Grass |
| Buildings | Building |
| Impervious Surface, Buildings | Impervious Surface |
| Roads | Roads |
| Trees | Trees |

Supplementary Table 2: Site Descriptions of Transects Pairs. Details of the characteristics of each transect pair, including descriptions of the vegetation, land use, and general surrounding area.

| Pair Name | Transect with water | Transect without water |
|---|---|---|
| 1 Barmston Drain South (Bridlington Avenue) | **Barmston Drain:** Major drain which runs through the city linearly, north to south, which connects to the River Hull. Riparian margins are well vegetated. Alongside the drain, there are strips of amenity grassland with trees. The section surveyed is surrounded by both residential and industrial urban areas. | **Central Ward:** Residential urban green spaces made up of amenity grasslands and mature trees. Fragmented green spaces are connected via grass verges. |
| 2 East Park | **East Park:** Large city park with woodlands, amenity grasslands and ornamental gardens. There is a large boating lake 1 km long, with multiple | **Rockford Fields:** Local nature reserve with a large patch of rough grassland. The site has a strip of mature trees with |

| | | |
|---|---|---|
| | islands. The site is bordered by residential urban areas on all sides. | scrub, and is surrounded by residential urban areas. |
| 3 Hessle Foreshore | **Hessle Foreshore:** Trail with scrub on one side and saltmarsh on the other, running alongside the shore of the Humber estuary. Some commercial and activity on the landward side towards the end of the section surveyed. (add details of reeds). | **Redcliff Road:** Urban area with many mature trees, sports fields and large gardens. Some commercial land use but mainly residential. |
| 4 Holderness Drain | **Holderness Drain:** Major drain which runs through the city linearly, north to south, which connects to the Humber estuary. Alongside the drain, there are strips of amenity grasslands. The section surveyed is surrounded by both residential and industrial urban areas. | **Withernsea Railway Track:** Disused railway line with scrub hedgerows and trees either side. Surrounding land includes sports fields, commercial and residential urban areas. Small drainage ditch mostly covered by vegetation present at the beginning of the surveyed section. |
| 5 Noddle Hill | **Noddle Hill:** Local Nature Reserve on the edge of the city which has been used as an example of passive rewilding. It has one large fishing pond and multiple nature ponds with substantial reedbeds. The rest of the area is made up of rough grassland, woodland and extensive scrub. Surrounded by farmland on three sides and amenity grassland on the other. | **Foredyke:** Infilled drain delineated by original hedgerows. The surrounding area is made up of amenity grassland with areas of rough grassland and woodland. Low density residential areas nearby with gardens and community green spaces. |
| 6 Barmston Drain North (Orchard Park) | **Barmston Drain:** Major drain which runs through the city linearly, north to south, connected to the River Hull. Riparian margins are well vegetated. Strips of amenity grasslands and scrub run alongside the drain. The section surveyed is surrounded by residential urban areas and farmland. | **Orchard Park:** Residential urban area with gardens and community green spaces made up of amenity grasslands. Patches of mature hedgerows and trees delineate open areas. |
| 7 River Hull (South) | **Oak road:** Transect along the river Hull. The section surveyed has soft edges, with extensive linear reedbeds. Commercial and residential urban areas nearby. | **Green Lane:** Muddy lane with overgrown hedges and mature trees, bordered by amenity grassland and gardens. Residential areas nearby. |
| 8 Pickering Park | **Pickering Park:** Large city park with woodlands, amenity grasslands and ornamental planting. There is a large lake with multiple islands. The site is bordered by residential urban areas on all sides. | **Boothferry Playing Fields:** Large amenity grassland with mature hedgerow running along the perimeter. Residential areas on all sides. |
| 9 River Hull (North) | **River Hull and Reservoir:** Transect along the river Hull. The section surveyed has soft edges (see above). The reservoir has hard edges with some self-seeded vegetation, including reedbeds. Residential and commercial urban areas nearby. | **Sutton Road:** Large amenity grassland with mature woodland and scrub along the edge and in patches, along a main road. Residential and commercial urban areas nearby. |

| | | |
|---|---|---|
| 10 Setting Dyke | **Garrowby Orchard and Setting Dyke:** Former amenity grassland which has been naturalised with unmown grasslands, tree planting and a section of daylighted drain. Rough grasslands and mature trees also present. Residential areas and farmland nearby. | **Dent Road:** Amenity grasslands with scrub and mature trees. Residential areas and farmland nearby. A train track runs along a large stretch of the surveyed area. |
| 11 The Deep | **The Deep:** Heavily modified estuarine habitat with hard edges. Mud flats exposed at low-tide. Commercial activity nearby including a public aquarium and its car park. Residential areas with gardens also present. | **Victoria Park:** Amenity grasslands with mature scrub and woodland running along the edges. Commercial and residential urban areas nearby. |

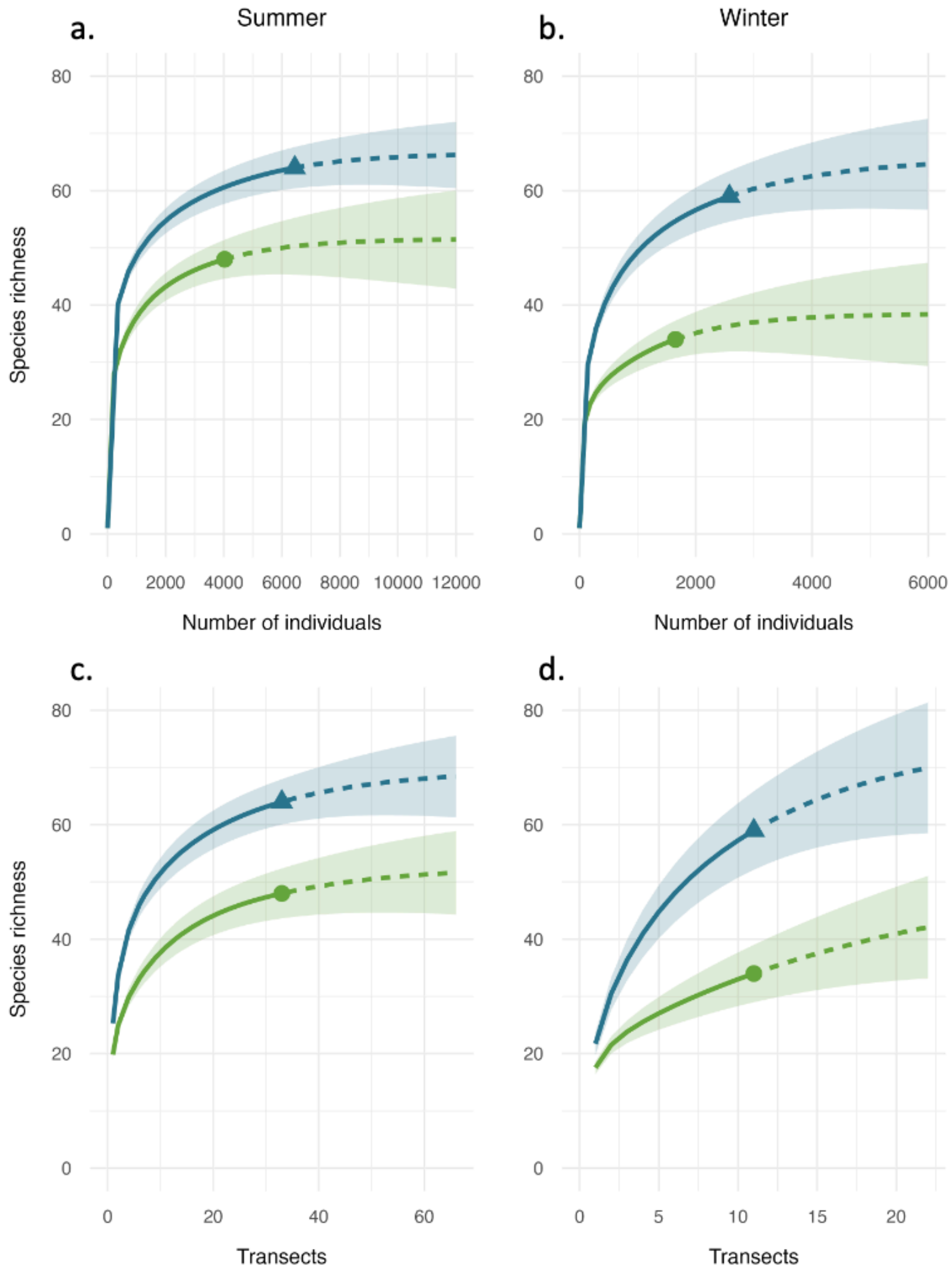


Supplementary Fig. 1. Species richness accumulation curves. Species accumulation is shown against bird abundance (a. and b.) and number of transects (c. and d.). Extrapolated estimates (dashed lines) were generated using the Chao1 estimator with 999 bootstrap permutations, extending to double the observed sampling effort. Transects with water are shown in blue with triangle symbols, transects without water are shown in green with circle symbols.

Supplementary Table 3: Analysis of variance for species richness and abundance in summer. Summary of species richness (a.) and abundance (b.) across the three summer recording periods (prior to data pooling). Variance was assessed using the aov function in R. Only the effect of water presence (yes/no) was statistically significant.

a. Bird Species Richness

| Effect | Df | Sum Sq | Mean Sq | F value | Pr(>F) |
|---|---|---|---|---|---|
| period | 2 | 5.7 | 2.9 | 0.121 | 0.886 |
| **water** | **1** | **491.8** | **491.8** | **20.755** | **2.67E-05** |
| period:water | 2 | 12.6 | 6.3 | 0.267 | 0.767 |
| Residuals | 59 | 1398.1 | 23.7 | | |

b. Bird Abundance

| Effect | Df | Sum Sq | Mean Sq | F value | Pr(>F) |
|---|---|---|---|---|---|
| period | 2 | 4383 | 2191 | 0.165 | 0.8486 |
| **water** | **1** | **88712** | **88712** | **6.663** | **0.0123** |
| period:water | 2 | 8577 | 4289 | 0.322 | 0.7259 |
| Residuals | 59 | 785487 | 13313 | | |

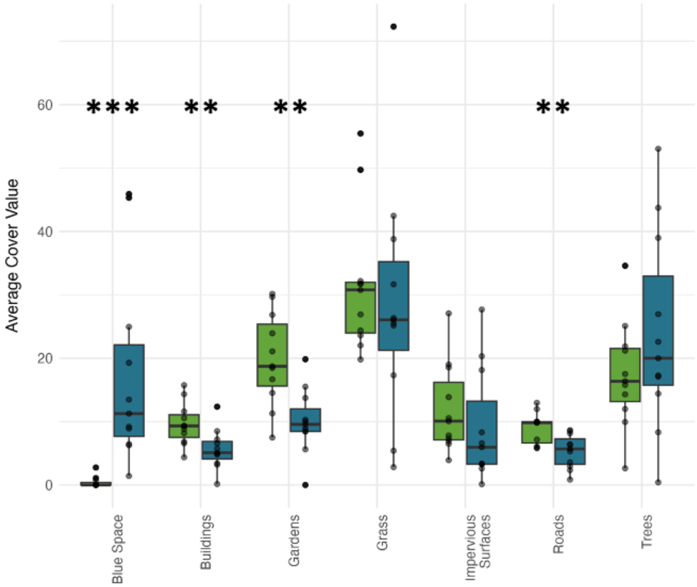


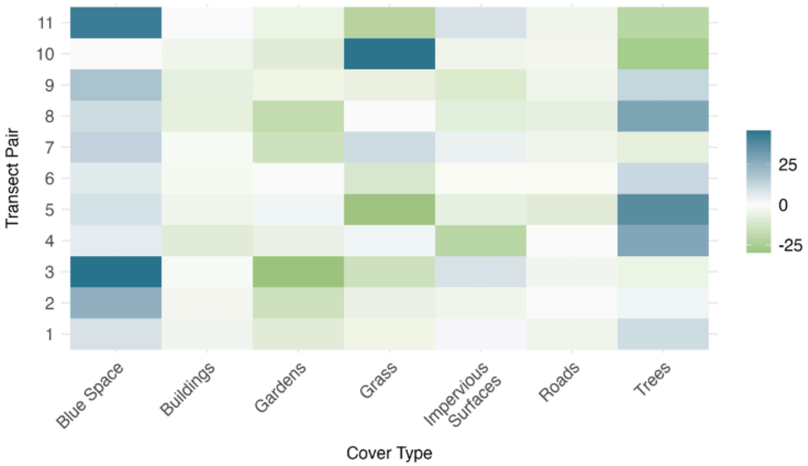


Supplementary Figure 2. Overview of land cover across transect pairs. On the left, comparisons of land cover (%) across transects with (blue) and without water (green). The most significant difference between them was blue space (W = 66, Z = –3.30, $p < 0.001$). Transects without blue space had significantly more garden cover (W = 2, Z = 2.98, $p = 0.003$), roads (W = 0, Z = 3.30, $p < 0.001$), and buildings (W = 1, Z = 3.10, adj. $p = 0.014$). On the right, pairwise differences in land cover across transects. Transects with water had more blue space at all sites (though this was minimal in Pair 10 due to a distant pond in the transect without water). Aside from blue space, tree cover and garden cover showed high variation, with garden cover generally higher in transects without water. Unnatural land cover types, buildings, impervious surfaces, and roads, were relatively balanced between transects.

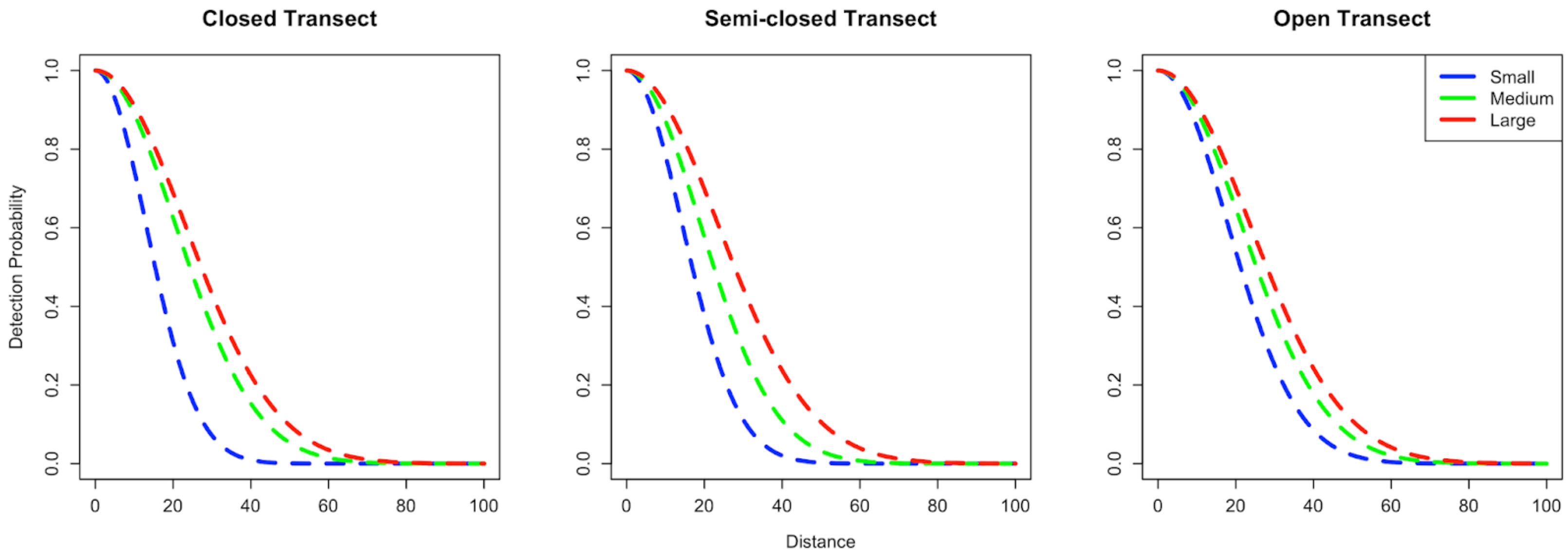


Supplementary Figure 3. Detection probability model. Detection decay over distance modelled with the two most explanatory covariates, bird size (size_class) and transect visibility (Region.Label), detection was modelled at 0.219 (SE = 0.003, CV = 0.015), indicating a good fit.

Supplementary Table 4. Distance decay model decision table. Summary of candidate models and covariates (a.) evaluated to identify the strongest predictors of bird detectability and the most accurate detection probability estimates (b.).

a.

| Model Rank | Detection Function | Model Formula | Detection Probability ($\hat{P_a}$) | Standard Error($\hat{P_a}$) | Change in AIC ($\Delta$AIC) |
|---|---|---|---|---|---|
| 1 | Half-normal | ~size_class * Region.Label | 0.219 | 0.003 | 0.000 |
| 2 | Half-normal | ~size_class * Region.Label * water | 0.218 | 0.003 | 4.484 |

| | | | | | |
|---|---|---|---|---|---|
| 3 | Half-normal | ~Region.Label + OBs + size_class | 0.219 | 0.003 | 5.221 |
| 4 | Half-normal | ~size_class + Region.Label | 0.220 | 0.003 | 6.174 |
| 5 | Half-normal | ~Region.Label + OBs + size_class + water | 0.219 | 0.003 | 6.712 |
| 6 | Half-normal | ~size_class + Region.Label + water | 0.220 | 0.003 | 7.738 |
| 7 | Half-normal | ~size_class + OBs + water | 0.220 | 0.003 | 17.995 |
| 8 | Half-normal | ~size_class + OBs | 0.221 | 0.003 | 22.625 |
| 9 | Half-normal | ~size_class | 0.221 | 0.003 | 24.142 |
| 10 | Half-normal | ~Region.Label + OBs | 0.228 | 0.003 | 192.118 |
| 11 | Half-normal | ~Region.Label + OBs + water | 0.228 | 0.003 | 193.610 |
| 12 | Half-normal | ~Region.Label | 0.228 | 0.003 | 196.783 |
| 13 | Half-normal | ~water | 0.229 | 0.003 | 213.968 |
| 14 | Half-normal | ~OBs | 0.229 | 0.003 | 214.726 |
| 15 | Half-normal | ~1 | 0.229 | 0.003 | 219.661 |

(Region.Label = transect visibility; OBs = human observer; size_class = bird size category; water = presence/absence of blue space (yes/no)

b.

| Bird Size | Region | Correction Factor |
|---|---|---|
| Small | Closed | 0.163 |
| Small | Semi-Closed | 0.258 |
| Small | Open | 0.289 |

| | | |
|---|---|---|
| Medium | Closed | 0.179 |
| Medium | Semi-Closed | 0.238 |
| Medium | Open | 0.295 |
| Large | Closed | 0.225 |
| Large | Semi-Closed | 0.269 |
| Large | Open | 0.297 |

Supplementary Table 5: Summary of Species Recorded. Table presenting key data on all recorded bird species (n = 79) during this study, including their common and scientific names, body mass, conservation status (BOCC5), habitat, and their presence across transects with and without water, during summer and winter surveys. Species unique to the transects with water are in bold.

| Common Name | Scientific Name | Mass (g) | BOCC5 Status | Habitat | % Presence in Transects with Water (Summer) | % Presence in Transects without Water (Summer) | % Presence in Transects with Water (Winter) | % Presence in Transects without Water (Winter) |
|---|---|---|---|---|---|---|---|---|
| Sparrowhawk | *Accipiter nisus* | 221 | Amber | Forest | 18.2 | 54.5 | 18.2 | 9.1 |
| **Sedge Warbler** | ***Acrocephalus schoenobaenus*** | 12 | Amber | Wetland | 36.4 | 0 | 0 | 0 |
| Reed Warbler | *Acrocephalus scirpaceus* | 12 | Green | Wetland | 54.5 | 9.1 | 0 | 0 |
| Long-tailed Tit | *Aegithalos caudatus* | 9 | Green | Forest | 63.6 | 81.8 | 45.5 | 72.7 |

| | | | | | | | | |
|---|---|---|---|---|---|---|---|---|
| Skylark | *Alauda arvensis* | 37 | Red | Grassland | 18.2 | 9.1 | 9.1 | 0 |
| **Kingfisher** | ***Alcedo atthis*** | 31 | Green | Riverine | 0 | 0 | 18.2 | 0 |
| Mallard | *Anas platyrhynchos* | 843 | Amber | Wetland | 90.9 | 27.3 | 72.7 | 9.1 |
| Greylag Goose | *Anser anser* | 3302 | Amber | Wetland | 45.5 | 9.1 | 54.5 | 9.1 |
| Pink-footed Goose | *Anser brachyrhynchus* | 2642 | Amber | Grassland | 0 | 0 | 9.1 | 9.1 |
| Swift | *Apus apus* | 38 | Red | Human Modified | 90.9 | 45.5 | 0 | 0 |
| Grey Heron | *Ardea cinerea* | 1443 | Green | Wetland | 18.2 | 9.1 | 18.2 | 0 |
| **Pochard** | ***Aythya ferina*** | 823 | Red | Wetland | 0 | 0 | 9.1 | 0 |
| **Tufted Duck** | ***Aythya fuligula*** | 701 | Green | Wetland | 18.2 | 0 | 27.3 | 0 |
| **Canada Goose** | ***Branta canadensis*** | 2812 | Green | Grassland | 18.2 | 0 | 27.3 | 0 |
| **Buzzard** | ***Buteo buteo*** | 759 | Green | Grassland | 18.2 | 0 | 0 | 0 |
| Goldfinch | *Carduelis carduelis* | 16 | Green | Woodland | 100 | 100 | 81.8 | 90.9 |
| **Treecreeper** | ***Certhia familiaris*** | 9 | Green | Forest | 9.1 | 0 | 0 | 0 |
| Cetti's Warbler | *Cettia cetti* | 13 | Green | Shrubland | 9.1 | 9.1 | 0 | 0 |
| Greenfinch | *Chloris chloris* | 26 | Red | Woodland | 72.7 | 90.9 | 18.2 | 9.1 |

| | | | | | | | | |
|---|---|---|---|---|---|---|---|---|
| Black-headed Gull | *Chroicocephalus ridibundus* | 284 | Amber | Wetland | 27.3 | 18.2 | 90.9 | 90.9 |
| **Marsh Harrier** | ***Circus aeruginosus*** | 704 | Amber | Wetland | 0 | 0 | 9.1 | 0 |
| Rock Dove | *Columba livia* | 354 | Green | Human Modified | 90.9 | 100 | 90.9 | 100 |
| Stock Dove | *Columba oenas* | 291 | Amber | Woodland | 54.5 | 9.1 | 18.2 | 18.2 |
| Woodpigeon | *Columba palumbus* | 490 | Amber | Woodland | 100 | 100 | 90.9 | 100 |
| Carrion Crow | *Corvus corone* | 570 | Green | Human Modified | 100 | 100 | 100 | 100 |
| Rook | *Corvus frugilegus* | 452 | Amber | Grassland | 9.1 | 18.2 | 0 | 0 |
| Jackdaw | *Corvus monedula* | 246 | Green | Human Modified | 18.2 | 27.3 | 18.2 | 18.2 |
| Whitethroat | *Curruca communis* | 15 | Amber | Shrubland | 72.7 | 36.4 | 0 | 0 |
| Lesser Whitethroat | *Curruca curruca* | 11 | Green | Shrubland | 9.1 | 9.1 | 0 | 0 |
| Blue Tit | *Cyanistes caeruleus* | 11 | Green | Forest | 100 | 100 | 81.8 | 100 |
| **Mute Swan** | ***Cygnus olor*** | 10682 | Green | Wetland | 45.5 | 0 | 36.4 | 0 |
| House Martin | *Delichon urbicum* | 15 | Red | Human Modified | 45.5 | 36.4 | 0 | 0 |

| | | | | | | | | |
|---|---|---|---|---|---|---|---|---|
| Great Spotted Woodpecker | *Dendrocopos major* | 75 | Green | Woodland | 36.4 | 27.3 | 18.2 | 0 |
| **Little Egret** | ***Egretta garzetta*** | 312 | Green | Wetland | 0 | 0 | 9.1 | 0 |
| **Reed Bunting** | ***Emberiza schoeniclus*** | 18 | Amber | Wetland | 36.4 | 0 | 9.1 | 0 |
| Robin | *Erithacus rubecula* | 18 | Green | Forest | 100 | 100 | 100 | 100 |
| Kestrel | *Falco tinnunculus* | 183 | Amber | Shrubland | 27.3 | 18.2 | 27.3 | 9.1 |
| Chaffinch | *Fringilla coelebs* | 24 | Green | Forest | 90.9 | 90.9 | 45.5 | 63.6 |
| Coot | *Fulica atra* | 717 | Green | Wetland | 54.5 | 9.1 | 54.5 | 0 |
| Moorhen | *Gallinula chloropus* | 340 | Amber | Wetland | 63.6 | 27.3 | 54.5 | 0 |
| Swallow | *Hirundo rustica* | 18 | Green | Human Modified | 81.8 | 18.2 | 0 | 0 |
| Herring Gull | *Larus argentatus* | 1091 | Red | Coastal | 100 | 100 | 100 | 90.9 |
| Common Gull | *Larus canus* | 413 | Amber | Coastal | 0 | 0 | 81.8 | 72.7 |
| Lesser Black-backed Gull | *Larus fuscus* | 762 | Amber | Coastal | 100 | 90.9 | 9.1 | 18.2 |
| Linnet | *Linaria cannabina* | 20 | Red | Shrubland | 54.5 | 9.1 | 0 | 0 |
| **Wigeon** | ***Mareca penelope*** | 770 | Amber | Wetland | 0 | 0 | 9.1 | 0 |
| **Gadwall** | ***Mareca strepera*** | 916 | Amber | Wetland | 18.2 | 0 | 9.1 | 0 |

| | | | | | | | | |
|---|---|---|---|---|---|---|---|---|
| **Goosander** | ***Mergus merganser*** | 1451 | Green | Riverine | 0 | 0 | 27.3 | 0 |
| Pied Wagtail | *Motacilla cinerea* | 24 | Amber | Human Modified | 18.2 | 18.2 | 0 | 9.1 |
| **Grey Wagtail** | ***Motacilla flava*** | 18 | Red | Grassland | 9.1 | 0 | 0 | 0 |
| **Curlew** | *Numenius arquata* | 803 | Red | Grassland | 0 | 0 | 9.1 | 0 |
| Great Tit | *Parus major* | 16 | Green | Woodland | 90.9 | 100 | 90.9 | 90.9 |
| House Sparrow | *Passer domesticus* | 27 | Red | Human Modified | 81.8 | 90.9 | 27.3 | 63.6 |
| Coal Tit | *Periparus ater* | 9 | Green | Forest | 18.2 | 45.5 | 18.2 | 9.1 |
| Cormorant | *Phalacrocorax carbo* | 2529 | Green | Wetland | 9.1 | 0 | 36.4 | 9.1 |
| **Pheasant** | ***Phasianus colchicus*** | 1120 | Green | Human Modified | 18.2 | 0 | 9.1 | 0 |
| Chiffchaff | *Phylloscopus collybita* | 8 | Green | Forest | 90.9 | 90.9 | 0 | 0 |
| Willow Warbler | *Phylloscopus trochilus* | 9 | Amber | Forest | 27.3 | 9.1 | 0 | 0 |
| Magpie | *Pica pica* | 218 | Green | Human Modified | 81.8 | 100 | 90.9 | 100 |

| | | | | | | | | |
|---|---|---|---|---|---|---|---|---|
| **Green Woodpecker** | ***Picus viridis*** | 176 | Green | Forest | 9.1 | 0 | 0 | 0 |
| **Great Crested Grebe** | ***Podiceps cristatus*** | 731 | Green | Wetland | 9.1 | 0 | 0 | 0 |
| Dunnock | *Prunella modularis* | 20 | Amber | Forest | 90.9 | 100 | 27.3 | 45.5 |
| **Ring-necked Parakeet** | ***Psittacula krameri*** | 116 | Green | Forest | 9.1 | 0 | 0 | 0 |
| Bullfinch | *Pyrrhula pyrrhula* | 24 | Amber | Forest | 18.2 | 27.3 | 18.2 | 9.1 |
| **Water Rail** | ***Rallus aquaticus*** | 111 | Green | Wetland | 0 | 0 | 9.1 | 0 |
| Goldcrest | *Regulus regulus* | 6 | Green | Forest | 18.2 | 18.2 | 9.1 | 0 |
| **Shoveler** | ***Spatula clypeata*** | 613 | Amber | Wetland | 0 | 0 | 9.1 | 0 |
| Siskin | *Spinus spinus* | 13 | Green | Forest | 0 | 0 | 27.3 | 27.3 |
| Collared Dove | *Streptopelia decaocto* | 149 | Green | Human Modified | 81.8 | 90.9 | 9.1 | 54.5 |
| Starling | *Sturnus vulgaris* | 77 | Red | Human Modified | 100 | 100 | 72.7 | 81.8 |
| Blackcap | *Sylvia atricapilla* | 17 | Green | Woodland | 100 | 90.9 | 0 | 0 |
| **Little Grebe** | ***Tachybaptus ruficollis*** | 169 | Green | Wetland | 9.1 | 0 | 9.1 | 0 |
| **Redshank** | ***Tringa totanus*** | 129 | Amber | Wetland | 0 | 0 | 18.2 | 0 |

| | | | | | | | | |
|---|---|---|---|---|---|---|---|---|
| Wren | *Troglodytes troglodytes* | 10 | Amber | Forest | 100 | 100 | 45.5 | 54.5 |
| **Redwing** | ***Turdus iliacus*** | 61 | Amber | Forest | 0 | 0 | 27.3 | 0 |
| Blackbird | *Turdus merula* | 103 | Green | Forest | 100 | 100 | 81.8 | 100 |
| Song Thrush | *Turdus philomelos* | 68 | Amber | Forest | 72.7 | 81.8 | 9.1 | 9.1 |
| **Mistle Thrush** | ***Turdus viscivorus*** | 117 | Red | Forest | 18.2 | 0 | 9.1 | 0 |
| **Lapwing** | ***Vanellus vanellus*** | 218 | Red | Wetland | 0 | 0 | 9.1 | 0 |

Supplementary Table 6: Bird species taxonomy and ecological characteristics. Table detailing the taxonomic details, trophic niche and size class assigned to all species. This information, as well as mass (see above) was used to calculate functional diversity.

| Common Name | Scientific Name | Genus | Family | Order | Trophic Niche | Primary Lifestyle | Size Class | Unique Trait Combination |
|---|---|---|---|---|---|---|---|---|
| Lapwing | *Vanellus vanellus* | Vanellus | Charadriidae | Charadriiformes | Omnivore | Terrestrial | medium | OmnivoreTerrestrialmedium |
| Mistle Thrush | *Turdus viscivorus* | Turdus | Turdidae | Passeriformes | Omnivore | Insessorial | medium | OmnivoreInsessorialmedium |
| Song Thrush | *Turdus philomelos* | Turdus | Turdidae | Passeriformes | Invertivore | Generalist | medium | InvertivoreGeneralistmedium |
| Blackbird | *Turdus merula* | Turdus | Turdidae | Passeriformes | Omnivore | Generalist | medium | OmnivoreGeneralistmedium |
| Redwing | *Turdus iliacus* | Turdus | Turdidae | Passeriformes | Invertivore | Insessorial | medium | InvertivoreInsessorialmedium |
| Wren | *Troglodytes troglodytes* | Troglodytes | Troglodytidae | Passeriformes | Invertivore | Insessorial | small | InvertivoreInsessorialsmall |

| | | | | | | | | |
|---|---|---|---|---|---|---|---|---|
| Redshank | *Tringa totanus* | Tringa | Scolopacidae | Charadriiformes | Aquatic predator | Terrestrial | medium | Aquatic predatorTerrestrialmedium |
| Little Grebe | *Tachybaptus ruficollis* | Tachybaptus | Podicipedidae | Podicipediformes | Aquatic predator | Aquatic | medium | Aquatic predatorAquaticmedium |
| Blackcap | *Sylvia atricapilla* | Sylvia | Sylviidae | Passeriformes | Omnivore | Insessorial | small | OmnivoreInsessorialsmall |
| Starling | *Sturnus vulgaris* | Sturnus | Sturnidae | Passeriformes | Omnivore | Insessorial | medium | OmnivoreInsessorialmedium |
| Collared Dove | *Streptopelia decaocto* | Streptopelia | Columbidae | Columbiformes | Omnivore | Terrestrial | medium | OmnivoreTerrestrialmedium |
| Siskin | *Spinus spinus* | Spinus | Fringillidae | Passeriformes | Granivore | Insessorial | small | GranivoreInsessorialsmall |
| Shoveler | *Spatula clypeata* | Spatula | Anatidae | Anseriformes | Aquatic predator | Aquatic | large | Aquatic predatorAquaticlarge |
| Goldcrest | *Regulus regulus* | Regulus | Regulidae | Passeriformes | Invertivore | Insessorial | small | InvertivoreInsessorialsmall |
| Water Rail | *Rallus aquaticus* | Rallus | Rallidae | Gruiformes | Aquatic predator | Terrestrial | medium | Aquatic predatorTerrestrialmedium |
| Bullfinch | *Pyrrhula pyrrhula* | Pyrrhula | Fringillidae | Passeriformes | Omnivore | Insessorial | small | OmnivoreInsessorialsmall |
| Ring-necked Parakeet | *Psittacula krameri* | Psittacula | Psittaculidae | Psittaciformes | Omnivore | Insessorial | medium | OmnivoreInsessorialmedium |
| Dunnock | *Prunella modularis* | Prunella | Prunellidae | Passeriformes | Invertivore | Terrestrial | small | InvertivoreTerrestrialsmall |
| Great Crested Grebe | *Podiceps cristatus* | Podiceps | Podicipedidae | Podicipediformes | Aquatic predator | Aquatic | large | Aquatic predatorAquaticlarge |
| Green Woodpecker | *Picus viridis* | Picus | Picidae | Piciformes | Invertivore | Terrestrial | medium | InvertivoreTerrestrialmedium |

| | | | | | | | | |
|---|---|---|---|---|---|---|---|---|
| Magpie | *Pica pica* | Pica | Corvidae | Passeriformes | Omnivore | Terrestrial | medium | OmnivoreTerrestrialmedium |
| Willow Warbler | *Phylloscopus trochilus* | Phylloscopus | Phylloscopidae | Passeriformes | Invertivore | Insessorial | small | InvertivoreInsessorialsmall |
| Chiffchaff | *Phylloscopus collybita* | Phylloscopus | Phylloscopidae | Passeriformes | Invertivore | Insessorial | small | InvertivoreInsessorialsmall |
| Pheasant | *Phasianus colchicus* | Phasianus | Phasianidae | Galliformes | Omnivore | Terrestrial | large | OmnivoreTerrestriallarge |
| Cormorant | *Phalacrocorax carbo* | Phalacrocorax | Phalacrocoracidae | Suliformes | Aquatic predator | Aquatic | large | Aquatic predatorAquaticlarge |
| Coal Tit | *Periparus ater* | Periparus | Paridae | Passeriformes | Invertivore | Insessorial | small | InvertivoreInsessorialsmall |
| House Sparrow | *Passer domesticus* | Passer | Passeridae | Passeriformes | Granivore | Terrestrial | small | GranivoreTerrestrialsmall |
| Great Tit | *Parus major* | Parus | Paridae | Passeriformes | Invertivore | Insessorial | small | InvertivoreInsessorialsmall |
| Curlew | *Numenius arquata* | Numenius | Scolopacidae | Charadriiformes | Aquatic predator | Terrestrial | large | Aquatic predatorTerrestriallarge |
| Grey Wagtail | *Motacilla flava* | Motacilla | Motacillidae | Passeriformes | Invertivore | Terrestrial | small | InvertivoreTerrestrialsmall |
| Pied Wagtail | *Motacilla cinerea* | Motacilla | Motacillidae | Passeriformes | Invertivore | Terrestrial | small | InvertivoreTerrestrialsmall |
| Goosander | *Mergus merganser* | Mergus | Anatidae | Anseriformes | Aquatic predator | Aquatic | large | Aquatic predatorAquaticlarge |
| Gadwall | *Mareca strepera* | Mareca | Anatidae | Anseriformes | Herbivore aquatic | Aquatic | large | Herbivore aquaticAquaticlarge |
| Wigeon | *Mareca penelope* | Mareca | Anatidae | Anseriformes | Omnivore | Terrestrial | large | OmnivoreTerrestriallarge |

| | | | | | | | | |
|---|---|---|---|---|---|---|---|---|
| Linnet | *Linaria cannabina* | Linaria | Fringillidae | Passeriformes | Granivore | Terrestrial | small | GranivoreTerrestrialsmall |
| Lesser Black-backed Gull | *Larus fuscus* | Larus | Laridae | Charadriiformes | Aquatic predator | Generalist | large | Aquatic predatorGeneralistlarge |
| Common Gull | *Larus canus* | Larus | Laridae | Charadriiformes | Omnivore | Terrestrial | large | OmnivoreTerrestriallarge |
| Herring Gull | *Larus argentatus* | Larus | Laridae | Charadriiformes | Aquatic predator | Generalist | large | Aquatic predatorGeneralistlarge |
| Swallow | *Hirundo rustica* | Hirundo | Hirundinidae | Passeriformes | Invertivore | Aerial | small | InvertivoreAerialsmall |
| Moorhen | *Gallinula chloropus* | Gallinula | Rallidae | Gruiformes | Omnivore | Terrestrial | medium | OmnivoreTerrestrialmedium |
| Coot | *Fulica atra* | Fulica | Rallidae | Gruiformes | Herbivore aquatic | Generalist | large | Herbivore aquaticGeneralistlarge |
| Chaffinch | *Fringilla coelebs* | Fringilla | Fringillidae | Passeriformes | Invertivore | Generalist | small | InvertivoreGeneralistsmall |
| Kestrel | *Falco tinnunculus* | Falco | Falconidae | Falconiformes | Vertivore | Aerial | medium | VertivoreAerialmedium |
| Robin | *Erithacus rubecula* | Erithacus | Muscicapidae | Passeriformes | Omnivore | Generalist | small | OmnivoreGeneralistsmall |
| Reed Bunting | *Emberiza schoeniclus* | Emberiza | Emberizidae | Passeriformes | Omnivore | Terrestrial | small | OmnivoreTerrestrialsmall |
| Little Egret | *Egretta garzetta* | Egretta | Ardeidae | Pelecaniformes | Aquatic predator | Terrestrial | medium | Aquatic predatorTerrestrialmedium |
| Great Spotted Woodpecker | *Dendrocopos major* | Dendrocopos | Picidae | Piciformes | Omnivore | Insessorial | medium | OmnivoreInsessorialmedium |

| | | | | | | | | |
|---|---|---|---|---|---|---|---|---|
| House Martin | *Delichon urbicum* | Delichon | Hirundinidae | Passeriformes | Invertivore | Aerial | small | InvertivoreAerialsmall |
| Mute Swan | *Cygnus olor* | Cygnus | Anatidae | Anseriformes | Herbivore aquatic | Aquatic | large | Herbivore aquaticAquaticlarge |
| Blue Tit | *Cyanistes caeruleus* | Cyanistes | Paridae | Passeriformes | Invertivore | Insessorial | small | InvertivoreInsessorialsmall |
| Lesser Whitethroat | *Curruca curruca* | Curruca | Sylviidae | Passeriformes | Invertivore | Insessorial | small | InvertivoreInsessorialsmall |
| Whitethroat | *Curruca communis* | Curruca | Sylviidae | Passeriformes | Invertivore | Insessorial | small | InvertivoreInsessorialsmall |
| Jackdaw | *Corvus monedula* | Corvus | Corvidae | Passeriformes | Omnivore | Terrestrial | medium | OmnivoreTerrestrialmedium |
| Rook | *Corvus frugilegus* | Corvus | Corvidae | Passeriformes | Omnivore | Terrestrial | large | OmnivoreTerrestriallarge |
| Carrion Crow | *Corvus corone* | Corvus | Corvidae | Passeriformes | Omnivore | Terrestrial | large | OmnivoreTerrestriallarge |
| Woodpigeon | *Columba palumbus* | Columba | Columbidae | Columbiformes | Omnivore | Terrestrial | large | OmnivoreTerrestriallarge |
| Stock Dove | *Columba oenas* | Columba | Columbidae | Columbiformes | Omnivore | Terrestrial | medium | OmnivoreTerrestrialmedium |
| Rock Dove | *Columba livia* | Columba | Columbidae | Columbiformes | Granivore | Terrestrial | medium | GranivoreTerrestrialmedium |
| Marsh Harrier | *Circus aeruginosus* | Circus | Accipitridae | Accipitriformes | Vertivore | Aerial | large | VertivoreAeriallarge |
| Black-headed Gull | *Chroicocephalus ridibundus* | Chroicocephal us | Laridae | Charadriiformes | Aquatic predator | Generalist | medium | Aquatic predatorGeneralistmedium |
| Greenfinch | *Chloris chloris* | Chloris | Fringillidae | Passeriformes | Granivore | Generalist | small | GranivoreGeneralistsmall |

| | | | | | | | | |
|---|---|---|---|---|---|---|---|---|
| Cetti's Warbler | *Cettia cetti* | Cettia | Scotocercidae | Passeriformes | Invertivore | Generalist | small | InvertivoreGeneralistsmall |
| Treecreeper | *Certhia familiaris* | Certhia | Certhiidae | Passeriformes | Invertivore | Insessorial | small | InvertivoreInsessorialsmall |
| Goldfinch | *Carduelis carduelis* | Carduelis | Fringillidae | Passeriformes | Granivore | Insessorial | small | GranivoreInsessorialsmall |
| Buzzard | *Buteo buteo* | Buteo | Accipitridae | Accipitriformes | Vertivore | Insessorial | large | VertivoreInsessoriallarge |
| Canada Goose | *Branta canadensis* | Branta | Anatidae | Anseriformes | Herbivore terrestrial | Terrestrial | large | Herbivore terrestrialTerrestriallarge |
| Tufted Duck | *Aythya fuligula* | Aythya | Anatidae | Anseriformes | Herbivore aquatic | Aquatic | large | Herbivore aquaticAquaticlarge |
| Pochard | *Aythya ferina* | Aythya | Anatidae | Anseriformes | Herbivore aquatic | Aquatic | large | Herbivore aquaticAquaticlarge |
| Grey Heron | *Ardea cinerea* | Ardea | Ardeidae | Pelecaniformes | Aquatic predator | Terrestrial | large | Aquatic predatorTerrestriallarge |
| Swift | *Apus apus* | Apus | Apodidae | Caprimulgiformes | Invertivore | Aerial | medium | InvertivoreAerialmedium |
| Pink-footed Goose | *Anser brachyrhynchus* | Anser | Anatidae | Anseriformes | Herbivore terrestrial | Terrestrial | large | Herbivore terrestrialTerrestriallarge |
| Graylag Goose | *Anser anser* | Anser | Anatidae | Anseriformes | Herbivore terrestrial | Terrestrial | large | Herbivore terrestrialTerrestriallarge |
| Mallard | *Anas platyrhynchos* | Anas | Anatidae | Anseriformes | Herbivore aquatic | Aquatic | large | Herbivore aquaticAquaticlarge |
| Kingfisher | *Alcedo atthis* | Alcedo | Alcedinidae | Coraciiformes | Aquatic predator | Insessorial | medium | Aquatic predatorInsessorialmedium |
| Skylark | *Alauda arvensis* | Alauda | Alaudidae | Passeriformes | Omnivore | Terrestrial | medium | OmnivoreTerrestrialmedium |

| | | | | | | | | |
|---|---|---|---|---|---|---|---|---|
| Long-tailed Tit | *Aegithalos caudatus* | Aegithalos | Aegithalidae | Passeriformes | Invertivore | Insessorial | small | InvertivoreInsessorialsmall |
| Reed Warbler | *Acrocephalus scirpaceus* | Acrocephalus | Acrocephalidae | Passeriformes | Invertivore | Insessorial | small | InvertivoreInsessorialsmall |
| Sedge Warbler | *Acrocephalus schoenobaenus* | Acrocephalus | Acrocephalidae | Passeriformes | Invertivore | Insessorial | small | InvertivoreInsessorialsmall |
| Sparrowhawk | *Accipiter nisus* | Accipiter | Accipitridae | Accipitriformes | Vertivore | Insessorial | medium | VertivoreInsessorialmedium |

Supplementary Table 7: Summary table of taxonomic and functional diversity tests. Results summary of pairwise comparisons between transects with and without water. Significant results are in bold.

| Metric | Season | Data | Test | Statistic (V) | P_Value |
|---|---|---|---|---|---|
| Shannon Index | Summer | Raw Counts | Wilcoxon Signed Rank | 46 | 0.278 |
| Simpson Index | Summer | Raw Counts | Wilcoxon Signed Rank | 37 | 0.765 |
| Pielou's Evenness | Summer | Raw Counts | Wilcoxon Signed Rank | 35 | 0.898 |
| **Total Species Richness** | **Summer** | **Raw Counts** | **Wilcoxon Signed Rank** | **51.5** | **0.016** |
| Shannon Index | Winter | Raw Counts | Wilcoxon Signed Rank | 35 | 0.898 |
| Simpson Index | Winter | Raw Counts | Wilcoxon Signed Rank | 30 | 0.831 |
| Pielou's Evenness | Winter | Raw Counts | Wilcoxon Signed Rank | 17 | 0.175 |
| Total Species Richness | Winter | Raw Counts | Wilcoxon Signed Rank | 54 | 0.068 |
| Shannon Index | Summer | Adjusted Counts | Wilcoxon Signed Rank | 45 | 0.32 |
| Simpson Index | Summer | Adjusted Counts | Wilcoxon Signed Rank | 35 | 0.898 |
| Pielou's Evenness | Summer | Adjusted Counts | Wilcoxon Signed Rank | 31 | 0.898 |

| | | | | | |
|---|---|---|---|---|---|
| **Total Species Richness** | **Summer** | **Adjusted Counts** | **Wilcoxon Signed Rank** | **64** | **0.007** |
| Shannon Index | Winter | Adjusted/Winter | Wilcoxon Signed Rank | 36 | 0.831 |
| Simpson Index | Winter | Adjusted/Winter | Wilcoxon Signed Rank | 30 | 0.831 |
| Pielou's Evenness | Winter | Adjusted/Winter | Wilcoxon Signed Rank | 14 | 0.102 |
| Total Species Richness | Winter | Adjusted/Winter | Wilcoxon Signed Rank | 40.5 | 0.201 |
| **AvTD** | **Summer** | **Raw Counts** | **Wilcoxon Signed Rank** | **57** | **0.032** |
| RaoQ | Summer | Raw Counts | Wilcoxon Signed Rank | 37 | 0.765 |
| Functional Redundancy | Summer | Raw Counts | Wilcoxon Signed Rank | 18 | 0.206 |
| **AvTD** | **Winter** | **Raw Counts** | **Wilcoxon Signed Rank** | **65** | **0.002** |
| RaoQ | Winter | Raw Counts | Wilcoxon Signed Rank | 38 | 0.7 |
| Functional Redundancy | Winter | Raw Counts | Wilcoxon Signed Rank | 21 | 0.32 |

Supplementary Table 8: Summary table of biomass comparisons. Comparisons of total bird mass in transects with and without blue space, using data with and without wetland species.

| Conservation Status | Season | Data | Test | Statistic (V) | P_Value |
|---|---|---|---|---|---|
| **Log Biomass (Kg)** | **Summer** | **Raw Counts** | **Wilcoxon Signed Rank** | **61** | **0.010** |
| **Log Biomass (Kg)** | **Winter** | **Raw Counts** | **Wilcoxon Signed Rank** | **59** | **0.010** |
| Log Biomass (Kg) | Summer | Raw Counts - Wetland birds | Wilcoxon Signed Rank | 45 | 0.320 |
| Log Biomass (Kg) | Winter | Raw Counts - Wetland birds | Wilcoxon Signed Rank | 42 | 0.465 |

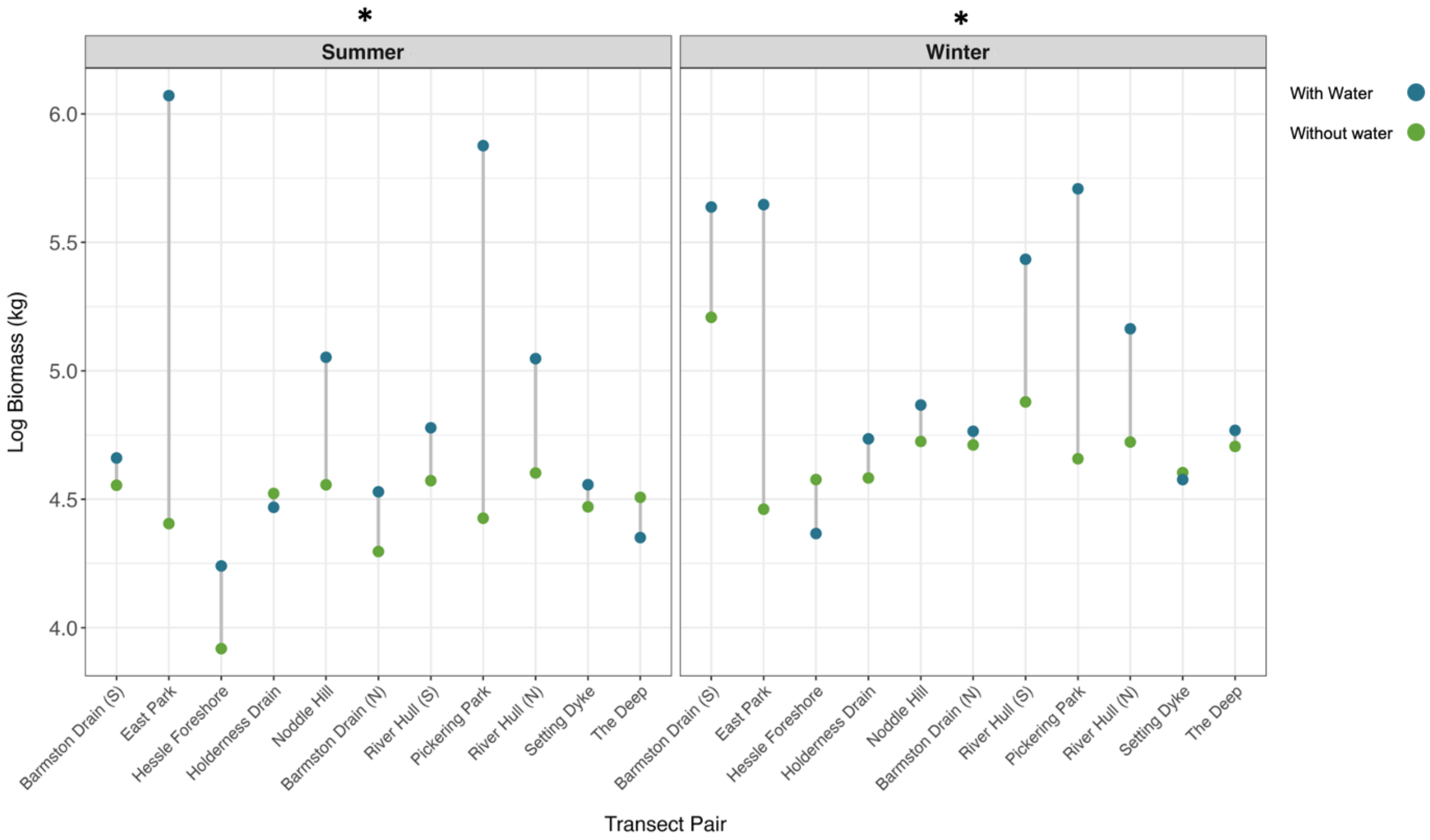


Supplementary Fig. 4. Log-transformed bird biomass (kg) across transect pairs during the Summer and Winter seasons. Each pair of points represents biomass in transects with (blue) and without (green) water, connected by lines to show the difference within sites. Biomass was significantly higher in transects with water during the Summer ($V = 61$, $p = 0.010$) and Winter ($V = 59$, $p = 0.019$), tested with paired Wilcoxon signed-rank tests.

Supplementary Table 9: Summary table of conservation status comparisons. Comparisons of the total number of conservation priority species in transects with and without water.

| Conservation Status | Season | Data | Test | Statistic (V) | P_Value |
|---|---|---|---|---|---|
| Green | Summer | Raw Counts | Wilcoxon Signed Rank | 45 | 0.081 |
| **Amber** | **Summer** | **Raw Counts** | **Wilcoxon Signed Rank** | **50.5** | **0.022** |
| **Red** | **Summer** | **Raw Counts** | **Wilcoxon Signed Rank** | **42** | **0.021** |
| Green | Winter | Raw Counts | Wilcoxon Signed Rank | 48 | 0.193 |
| **Amber** | **Winter** | **Raw Counts** | **Wilcoxon Signed Rank** | **60** | **0.017** |
| Red | Winter | Raw Counts | Wilcoxon Signed Rank | 26.5 | 0.668 |
